# Low CAR affinity enhances T-cell synapse dynamics and activation

**DOI:** 10.64898/2026.10.09.757739

**Authors:** Chikim Nguyen, Gerard Castro-Linares, Lucrezia C.D.E. Gatti, Zsolt Sebestyen, Ricardo A Fernandes, Sander J. Tans, Megan V. Farrell, Kristina A. Ganzinger

## Abstract

Low affinity chimeric antigen receptors (CARs) can clinically outperform high-affinity variants, yet the molecular mechanisms underlying affinity-dependent activity remain unclear. Here, we show that low CAR affinity enables faster CAR cluster motion, which is central to T-cell activation. We compared low-affinity CAT and high-affinity FMC63 CARs signaling at different stages of activation on supported lipid bilayers functionalized with defined CD19 densities. At low CAR expression on dense CD19 bilayers, low-affinity CAT CARs formed more mature and dynamic synapses, recruited ZAP70 more efficiently, and showed faster microcluster motion. These proximal signaling changes were accompanied by stronger calcium flux and increased NFAT nuclear translocation. Our data support a model in which weaker, lower-affinity CAR-antigen interactions promote receptor turnover, microcluster mobility, and signaling-adaptor recruitment, thereby enhancing downstream signal propagation. Our work provides a mechanistic basis for the superior functional performance reported for affinity-tuned CD19 CAR-T cells.

## Main

Chimeric antigen receptors (CARs) are synthetic immunoreceptors that combine the specificity of monoclonal antibodies with the signaling machinery of the T cell antigen receptor (TCR-CD3) and are used for cancer immunotherapy^1,2^. A typical CAR comprises an antibody-derived scFv specific for tumor antigens (e.g., CD19, BCMA or CD22), a co-stimulatory domain (e.g., 4-1BB or CD28) and the CD3*ζ* signaling module that together drive antigen-dependent T cell activation. Among the seven FDA-approved CAR-T cell products, five CARs target CD19 and have demonstrated remarkable efficacy in relapsed or refractory B-cell lymphomas and acute lymphoblastic leukemia (ALL)^1,3^. Notably, all five CD19-directed CARs employ the high-affinity FMC63 scFv as their antigen-binding domain, while multiple studies have now shown that CARs with a lower-affinity CAT scFV domain (CAT CAR) display stronger antigen-specific cytotoxicity, less toxicity and improved antitumor activity in both in vitro and in vivo preclinical models compared to high-affinity FMC63 CARs^4–6^. However, how different antigen-binding domains engineered in CAR constructs, including CAT and FMC63, influence the early CAR-T cell activation that shapes these distinct phenotypes and outcomes remains poorly understood.

Addressing these questions requires a quantitative insight into how CAR designs affect immune synapse formation and initiate intracellular signaling. To this end, we established supported-lipid bilayer (SLB)-based model system of tumor cell membranes, which enabled high-resolution imaging of CAR spatiotemporal organization during early CAR-T cell activation. This approach revealed how CAR design affects CAR microclusters formation and CD3*ζ* -binding signaling protein ZAP70 (Zeta-chain-associated protein kinase 70) recruitment, synapse organization and downstream signaling responses, all previously linked to clinical outcomes in patients^7^. We analyzed CAR immune synapse maturation based on convolutional neural network (CNN) classification and showed that the spatiotemporal organization and dynamics of CAR microclusters binding to CD19 and recruiting ZAP70 (CD19-CAR/ZAP70 microclusters) was influenced by antigen density, CAR expression, and CAR-antigen affinity. Low-affinity CAT CAR-T cells formed more active and mobile synapses than the high-affinity FMC63 CAR-T cells, recruiting ZAP70 more efficiently and triggering stronger downstream signaling, which may help explain their superior clinical performance.

## Results

### CAR affinity and CD19 density shape the spatiotemporal organization of CD19-CAR/ZAP70 microclusters and immune synapse formation

Quantifying receptor-antigen interactions and downstream signaling in T cell-tumor cell conjugates is hindered by 3D cell movement, phototoxicity from intense excitation, noise from scattering and cellular fluorescence. To overcome this, we used a planar glass-SLB as a surrogate target cell surface for the formation of immune synapses with CAR-T cells (Jurkat E6.1 cells expressing CARs). This allows for total internal reflection fluorescence microscopy (TIRF) microscopy and subsequent single-molecule analysis of plasma membrane-proximal events (Figure 1a). Our SLBs incorporated 18:1 DGS-NTA(Ni), allowing precise attachment of polyhistidine-tagged recombinant proteins, including the CAR ligand CD19 and the adhesion receptor lymphocyte function-associated antigen 1 (LFA-1) ligand ICAM-1. Given Jurkat E6.1 cells express low levels of LFA-1^8^, limiting ICAM-1-mediated attachment, MCC-DOPE lipid was added to the SLB to allow functionalization with thiol-modified synthetic DNA. A complementary DNA sequence with a cholesterol moiety was added to the cells to enhance cell adhesion. Incorporation of the DNA adhesion system (modeled on the intercellular spacing of LFA-1/ICAM-1^9^) increased the cell-SLB contact footprint by approximately 2.5-fold (Figure S1a-c) and allowed us to titrate CAR antigen designs down to roughly 1 molecule/µm^2^.

**Figure 1.**
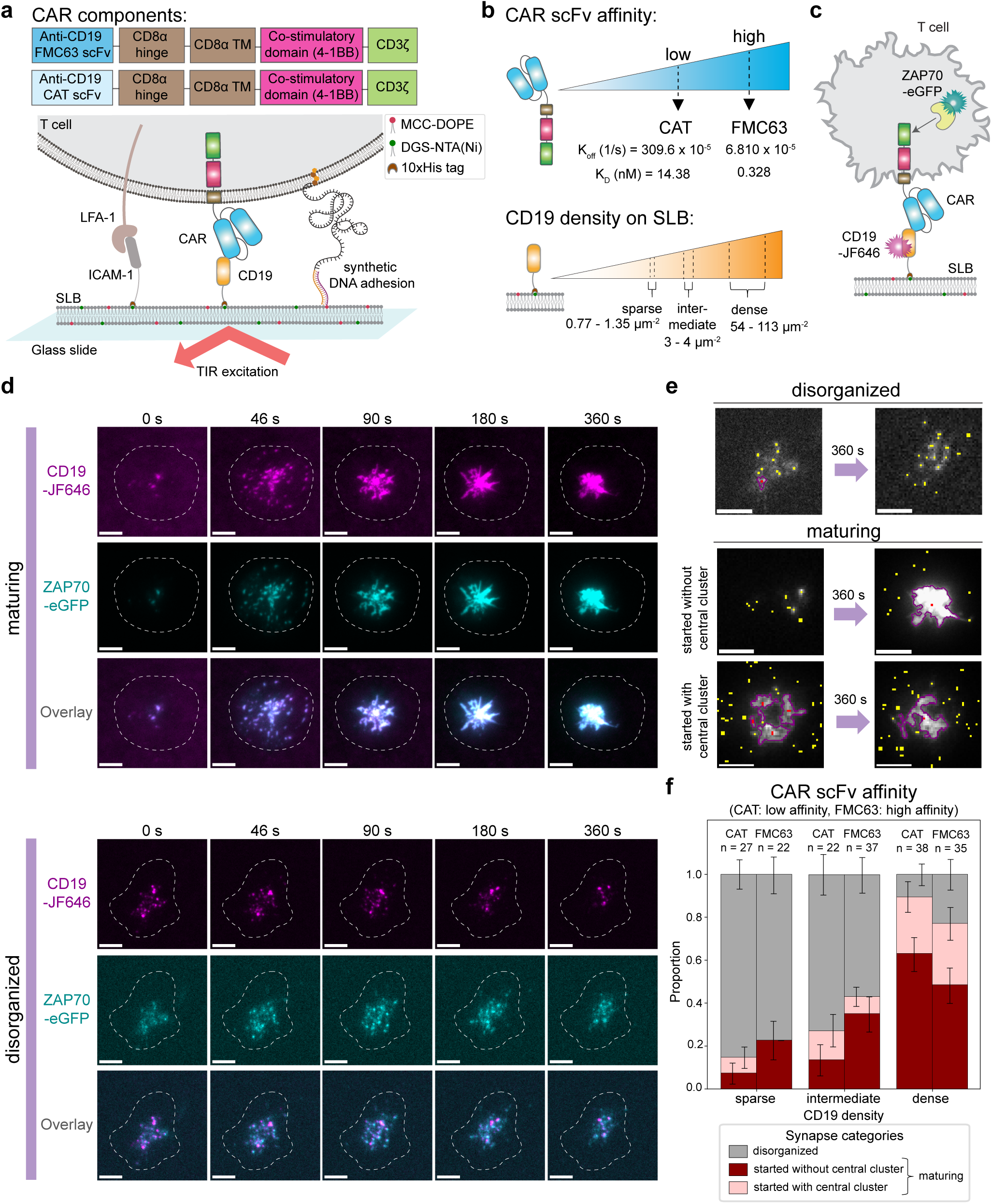
Increasing CD19 density led to more mature synapse formation for both low- and high-affinity CARs with low CAR expression. (a) Schematic of the CD19-targeting CAR construct and SLB platform used to investigate immune synapse formation and early CAR signaling. The SLB displays CD19 and ICAM-1 to engage CAR and LFA-1 on T cells, respectively, while a DNA-mediated adhesion system enhances cell-bilayer contact. CD19 and ICAM-1 are anchored through 10×His-tag binding to 18:1 DGS-NTA(Ni) lipids, and thiol-modified single-stranded DNA is coupled to MCC-DOPE lipids. (b) Schematic of the experimental conditions used in this study, including CD19-targeting CARs with low-affinity (CAT; *k*_off_ = 309.6 × 10*^−^*^5^ s*^−^*^1^, *K_D_* = 14.38 nM^4^) or high-affinity (FMC63; *k*_off_ = 6.810 × 10*^−^*^5^ s*^−^*^1^, *K_D_* = 0.328 nM^4^) scFvs and three CD19 density regimes on SLBs: sparse (0.77-1.35 molecules/µm^2^), intermediate (3-4 molecules/µm^2^), and dense (54-113 molecules/µm^2^). **(c)** Schematic illustrating the spatiotemporal recruitment of ZAP70-eGFP to the CAR following engagement with CD19-JF646, enabling visualization and quantification of CD19-CAR/ZAP70 colocalization complexes. **(d)** Representative time-lapse TIRF image sequences showing the spatiotemporal recruitment of ZAP70-eGFP (cyan) following CAR engagement with CD19-JF646 (magenta) in two cells exhibiting either mature or disorganized immune synapses. Dashed outlines indicate individual cell footprints on SLBs, manually determined from maximum-intensity projections of the CD19 signal over the imaging period. Scale bars are 5 µm. **(e)** Time-lapse sequences of ZAP70-eGFP recruitment to CD19-CAR microclusters were used in CNN-based approach to classify CAR immune synapses as either disorganized or maturing, and whether the maturing synapse started with or without an initial central cluster(s). Representative TIRF images acquired at the onset of imaging and 360 s later show the temporal evolution of CD19-CAR/ZAP70 organization in each synapse category. Small peripheral clusters are highlighted in yellow, large clusters are outlined in magenta with red dots indicate cluster centroids. Scale bars are 5 µm. **(f)** Stacked bar plots showing the proportion of low-CAR-expressing cells classified into different synapse maturation categories at sparse, intermediate, and dense CD19 densities on SLBs for low-affinity (CAT) and high-affinity (FMC63) CAR-T cells. Error bars represent the standard errors of the binomial proportions. Numbers of analyzed cells were: CAT CAR-T cells, *n* = 27 (sparse), *n* = 22 (intermediate), and *n* = 38 (dense); FMC63 CAR-T cells, *n* = 22 (sparse), *n* = 37 (intermediate), and *n* = 35 (dense).

To determine how CAR affinity shapes early CAR signaling events, previously shown to include similar adaptors to native T cell receptors (TCRs)^10,11^, we engineered Jurkat E6.1 T cells to express either low-affinity (CAT, K_D_ = 14.38 nM^4^) or high-affinity (FMC63, K_D_ = 0.328 nM^4^) CD19-targeting 4-1BB-CD3*ζ* CARs. We then assessed ligand engagement using CD19 labeled with Janelia Fluor 646 (JF646), receptor triggering through ZAP70-eGFP recruitment to phosphorylated CD3*ζ* tails, as well as immune synapse formation by following the clustering of CD19-CAR/ZAP70 complexes (Figure 1a-c). To modulate ligand availability, a key determinant of CAR-T cell activity^12^, we tuned the surface density of CD19 on SLBs to create either sparse (0.77-1.35 molecules/µm^2^), intermediate (3-4 molecules/µm^2^), or dense (54-113 molecules/µm^2^) conditions (Figure 1b), matching the cell surface expression level on the pre-B acute lymphoblastic leukemia cell line NALM6/WT (27,000 CD19 molecules, ∼86 CD19 molecules/µm^2^), NALM6/12-39 (2,000 CD19 molecules, ∼6.4 CD19 molecules/µm^2^) or NALM6/12-4 (200 CD19 molecules, ∼0.64 CD19 molecules/µm^2^), respectively^6^. Moreover, to better resolve distinct CAR signaling behaviors, we stratified our CAR-expressing T cells (Figure S2a-c) into low- and high-CAR-expressing groups.

Imaging immune synapse formation between CAR-T cells and CD19-SLBs indicated the formation of CD19-CAR microclusters at these interfaces that robustly recruited ZAP70, consistent with previous studies^10,13,14^ (Figure 1d) while the DNA adhesion system on its own did not induce CAR clustering or any unspecific ZAP70 recruitment in the absence of CD19 (Figure S1d,e). Notably, our data revealed two distinct spatiotemporal patterns of CD19-CAR clusters recruiting ZAP70 that we refer to as maturing and disorganized immune synapses. A maturing synapse is characterized by the formation of peripheral signaling microclusters that undergo centripetal movement and ultimately coalesce into a compact central supramolecular activation cluster (cSMAC), forming the canonical “bull’s eye” architecture well known from TCR signaling^15–18^. However, CAR synapses do not always follow this maturation trajectory. In these disorganized synapses, ZAP70 is recruited to and colocalizes with CD19-CAR microclusters, but these microclusters fail to merge into a centralized cSMAC and instead persist as small, dispersed clusters, consistent with recent studies reporting similar patterns of CAR synapse organization^14^. For further analysis of the immune synapses across antigen densities and CAR constructs, cells that formed a mature synapse during the 360 to 400 s imaging window were divided into two subgroups based on whether the large central CD19-CAR/ZAP70 cluster(s) were already present at the start of imaging, or not (Figure 1e). In both cases, we observed peripheral microclusters that migrated inward and ultimately fused with the central structure. To achieve an unbiased and quantitative classification of CAR-T cell immune synapse maturation, we implemented a CNN-based classifier trained on TIRF data with ZAP70 recruitment dynamics (Figure S3). Notably, most synapses classified as “maturing” were categorized as “started without central cluster”, indicating that we captured the majority of synapses from the onset of their maturation (Figure 1f). In contrast, synapses classified as “started with central cluster” were typically imaged after the central cluster had already formed and exhibited little to no subsequent dynamics in the peripheral microclusters. Consequently, these synapses were excluded from further analyses.

In low-CAR-expressing T cells, the relative proportions of mature versus disorganized synapses strongly depended on CD19 density: increasing CD19 density from sparse to intermediate conditions led to around 2-fold increases in mature synapse formation for both low- and high-affinity CARs (Figure 1f). A similar trend was observed in high-CAR-expressing T cells (Figure S4). While the overall proportion of mature synapses was higher in high-than in low-CAR-expressing cells (Figure 1f, Figure S4), CAR expression had only a mild effect on synapse maturation at dense CD19 densities, where mature synapse formation was uniformly elevated across all conditions (77% to 94%). Interestingly, among low-CAR-expressing cells, those with low-affinity CAR formed mature synapses more frequently than their high-affinity counterparts (89% versus 77%) (Figure 1f). Together, our results indicate that the formation of mature CAR immune synapses are jointly shaped by CD19 antigen density and CAR expression level on the engineered T cells, with increases in either parameter promoting the formation of mature synapses.

### Enhanced ZAP70 recruitment efficiency to CARs in maturing synapses

To further understand the differences between disorganized and maturing synapses that started without central cluster, we next quantified CD19-CAR engagement and ZAP70 recruitment by integrating the fluorescence intensities of all CD19-JF646 and ZAP70-eGFP clusters detected, including both the larger central clusters and the smaller peripheral microclusters, at each synapse and averaging these values across frames for each cell (Figure 2a). Compared with disorganized synapses, maturing synapses exhibited similar CD19 binding (Figure 2b), but higher ZAP70 recruitment (Figure 2c). As a result, the ratio of ZAP70 recruited per CD19 molecule is significantly higher in maturing compared to disorganized synapses to dense CD19 surfaces (for both low- and high-affinity CARs). Thus, CAR-T cells that form mature immune synapses have a higher efficiency of activation and ZAP70 recruitment than cells that form disorganized synapses (Figure 2d). Given that most high-CAR-expressing cells interacting with dense CD19 surfaces formed mature synapses (92-94%, Figure S4), the number of disorganized cells under this condition was too low to allow a direct comparison with low-CAR-expressing cells on dense CD19 surfaces (Figure S5). Among high-CAR-expressing cells interacting with intermediate CD19 surfaces, cells forming mature synapses exhibited both greater CD19 binding and ZAP70 recruitment compared with cells forming disorganized synapses (Figure S5). However, the resulting ZAP70-to-CD19 ratio did not differ between them, suggesting that at high receptor expression levels, differences in ZAP70 recruitment efficiency relative to CD19 engagement are largely diminished.

**Figure 2.**
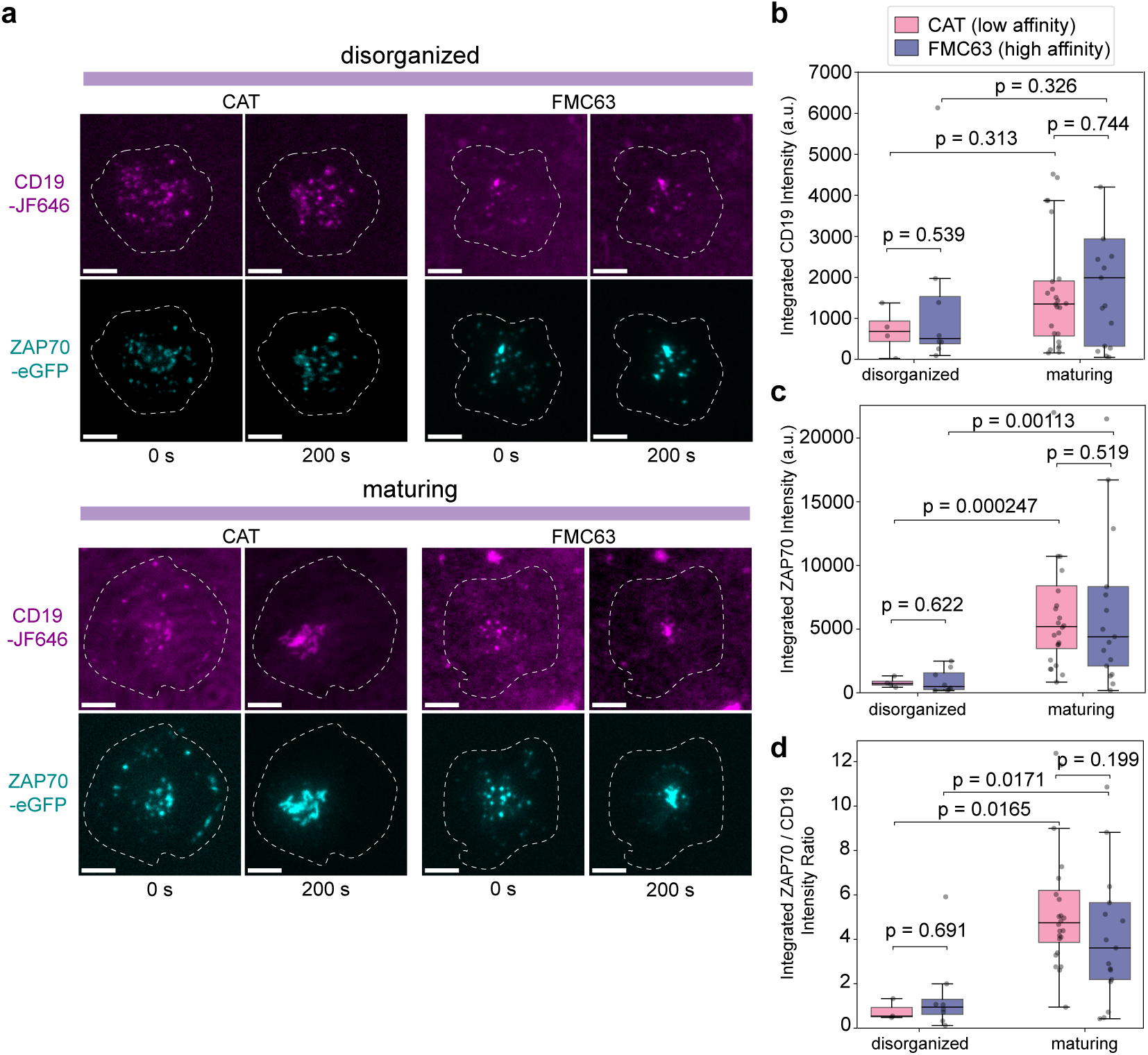
Maturing synapses showed similar CD19 engagement but increased ZAP70 recruitment compared with disorganized synapses, indicating more efficient signaling. **(a)** Representative TIRF images of low-affinity (CAT) and high-affinity (FMC63) CAR-T cells acquired at the onset of imaging (t = 0 s) and 200 s later, illustrating the temporal evolution of CD19-CAR (magenta)/ZAP70 (cyan) organization into either disorganized or maturing synapses. Dashed outlines indicate individual cell footprints on SLBs, manually determined from maximum-intensity projections of the CD19 signal over the imaging period. Scale bars are 5 µm. Box plot of the per-cell integrated **(b)** CD19-JF646 intensity, **(c)** ZAP70-eGFP intensity, and (d) ZAP70/CD19 intensity ratio for all CAR clusters detected during time-lapse TIRF imaging of disorganized and maturing synapses starting with a central cluster in low-affinity (CAT) and high-affinity (FMC63) CAR-T cells expressing low CAR levels on dense CD19 SLBs. Boxes indicate the median and interquartile range, and individual points represent a single cell. Numbers of analyzed cells were: CAT CAR-T cells, *n* = 4 (disorganized) and *n* = 24 (maturing); FMC63 CAR-T cells, *n* = 8 (disorganized) and *n* = 17 (maturing).

### Low-affinity CARs show more efficient CD19 binding and ZAP70 recruitment

To further dissect early activation differences between CARs of different affinities, we next quantified ZAP70 recruitment in small, early activation microclusters and followed their dynamics in maturing cells that started with no central cluster. These small CD19-CAR/ZAP70 clusters are thought to serve as the major early signaling hubs and display minimal size variability, allowing for more reliable quantification^19^. Conversely, large central clusters generally mark terminal stages of signaling, where activated components accumulate before being internalized through endocytosis^20^ or being released as exosomes^21^ or ectosomes^22–24^ for receptor recycling and eventual degradation and were therefore not included in our subsequent analyses.

By independently tracking the trajectory of small CD19 and ZAP70 clusters and then overlaying their tracks, we could follow CD19 microcluster dynamics before and during ZAP70 recruitment to CARs and thus, pinpoint the moment when ZAP70 first engaged each CD19-CAR cluster (Figure 3a,b). Under sparse CD19 conditions, high-affinity CARs bound slightly less CD19 compared with low-affinity CARs (0.9-fold lower during ZAP70 colocalization; Figure 3b,c and Figure S8a), but recruited similar amounts of ZAP70 (Figure 3b,d and Figure S8b), resulting in a modest increase in ZAP70-per-CD19 efficiency (1.1-fold during ZAP70 colocalization; Figure 3e, Figure S6 and Figure S8c). Increasing CD19 density to intermediate and dense conditions enabled high-affinity CARs at low expression to bind more CD19 (1.2-fold higher during ZAP70 colocalization) than low-affinity CARs, both before and during ZAP70 colocalization (Figure 3b,c and Figure S8a). This indicates that high-affinity CARs could exploit increased CD19 availability, whereas low-affinity CARs, constrained by faster off-rates, were unable to achieve comparable binding to the high-affinity CAR at low receptor abundance. Despite this, low-affinity CARs recruited similar or higher amounts of ZAP70 than high-affinity CARs at intermediate and dense CD19 densities, respectively (Figure 3b,d and Figure S8b). As a result, their ZAP70-per-CD19 efficiency surpassed that of high-affinity CARs at both intermediate and dense densities (1.2-fold and 1.9-fold higher, respectively) (Figure 3e, Figure S6 and Figure S8c). This suggests that while high-affinity CARs signal stronger at very low antigen densities, the low-affinity CARs outperform the high-affinity CARs at higher antigen densities.

**Figure 3.**
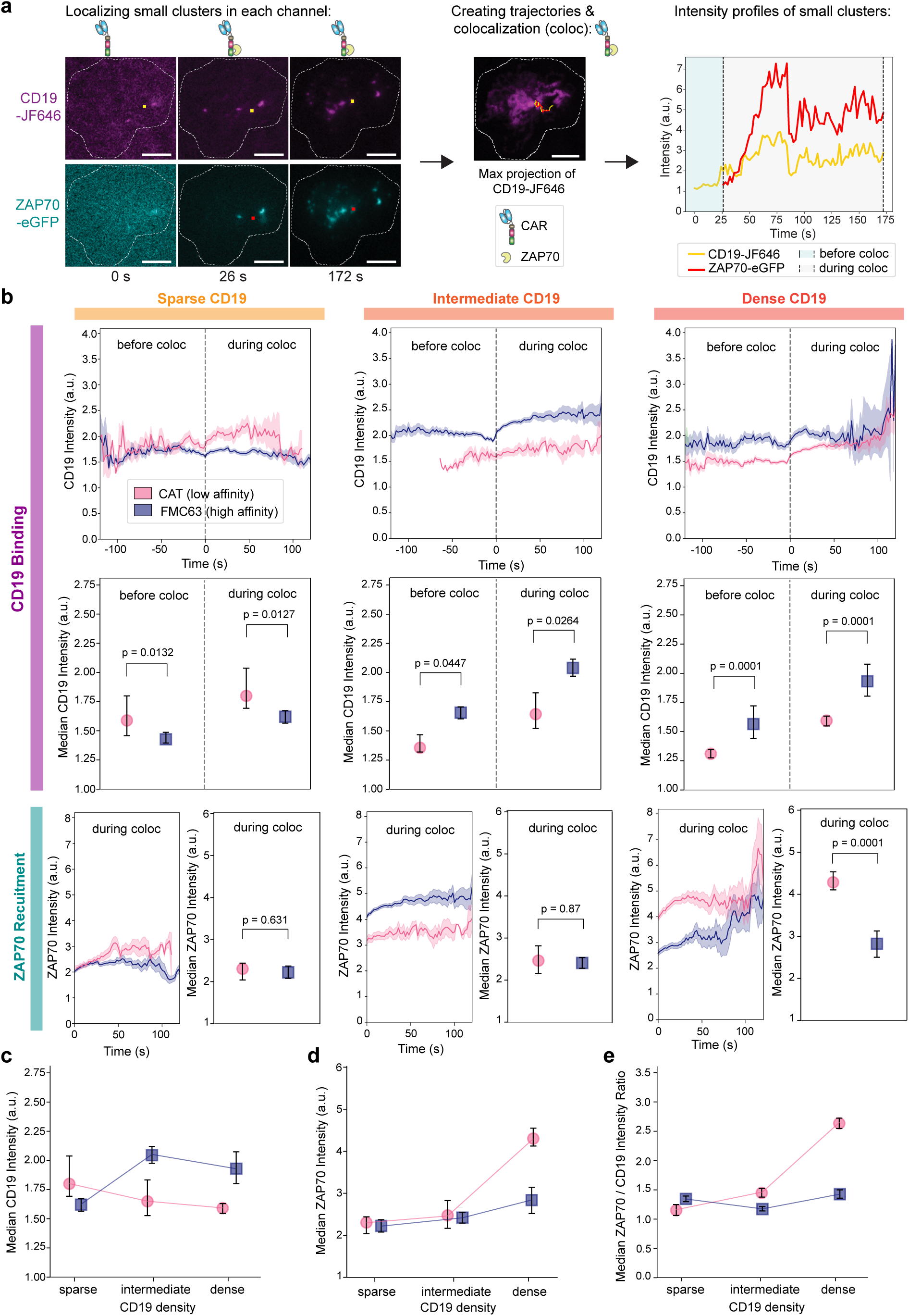
Low-affinity CARs showed more efficient CD19 binding and ZAP70 recruitment. **(a)** Representative time-lapse TIRF images acquired at t = 0 s, 26 s (ZAP70 recruitment), and 200 s, showing the detection and tracking of CD19-JF646 (yellow) and ZAP70-eGFP microclusters (red) in separate imaging channels. Microcluster trajectories were reconstructed from localized detections across frames, and CD19-CAR/ZAP70 colocalization events were identified by trajectory overlap. Fluorescence intensity traces of individual trajectories reveal the onset of ZAP70 recruitment and the associated temporal changes in CD19 and ZAP70 signals. Dashed outlines indicate individual cell footprints on SLBs, manually determined from maximum-intensity projections of the CD19 signal over the imaging period. Scale bars are 5 µm. **(b)** CD19-JF646 and ZAP70-eGFP fluorescence intensity traces, shown as geometric means with SEM estimated from log-transformed values, from small CD19-CAR and ZAP70 microcluster trajectories, together with dot plots showing the corresponding median fluorescence intensities with bootstrap confidence intervals, generated by low-affinity (CAT) and high-affinity (FMC63) CAR-T cells expressing low CAR levels interacting with sparse, intermediate, or dense CD19 SLBs. **(c)** Summary of CD19 engagement to CAR during ZAP70 colocalization. Data correspond to the median CD19-JF646 intensities shown in (b). **(d)** Summary plot showing enhanced ZAP70 recruitment to CD19-CAR microclusters with increasing CD19 density on SLBs, particularly for low-affinity CARs. Data correspond to the median ZAP70-eGFP intensities shown in (b). **(e)** Summary plot showing increased ZAP70 recruitment efficiency relative to CD19-CAR engagement with increasing CD19 density on SLBs, particularly for low-affinity CARs. Data correspond to the median ZAP70-eGFP/CD19-JF646 intensity ratios shown in Figure S6. Number of analyzed tracks under different CD19 densities: sparse CD19, CAT (n = 55) and FMC63 (n = 218); intermediate CD19, CAT (n = 29) and FMC63 (n = 784); dense CD19, CAT (n = 447) and FMC63 (n = 147).

Interestingly, in high-CAR-expressing cells with mature synapses that started with no central cluster, low-affinity CARs bound similar or larger amounts of CD19 than high-affinity CARs at intermediate or dense CD19 densities, particularly during ZAP70 colocalization (Figure S7a and Figure S8a). At low CAR expression levels, ZAP70 recruitment per CD19 was higher for low-compared to high-affinity CARs at intermediate and dense CD19 conditions (Figure S6b,c and Figure S8b,c), indicating more efficient signaling adaptor recruitment per engaged ligand. This observation is consistent with the possibility that the faster dissociation rate of the low-affinity CAR permits serial engagement of multiple CD19 molecules, thereby increasing the number of signaling-competent CARs generated per ligand. In contrast, this difference was largely lost at high CAR expression levels, where high-affinity FMC63 CARs exhibited similar, and in dense CD19 conditions, greater ZAP70 recruitment per CD19 molecule, though less CD19 binding overall (Figure S7b,c and Figure S8b,c). This may indicate that at high CAR expression and under dense ligand conditions, the high-affinity receptors are more constrained in dense receptor clusters, remaining tightly bound to individual CD19 molecules (Figure S7a) and limiting ligand turnover and reducing opportunities for additional CD19 engagement. In contrast, for low-affinity CAT CARs, increasing the CAR density (at high CAR expression levels) was associated with the formation of larger microclusters and greater CD19 accumulation. However, the increased size and packing of these microclusters may reduce the accessibility of ZAP70 binding sites, resulting in a similar amount of ZAP70 recruited per microcluster despite the greater CD19 binding and an overall reduction in Zap70/CD19 ratio (Figure S7a, b, c and Figure S8a, b, c). Together, these results indicate that CD19 density, CAR affinity and CAR expression level jointly determine both ligand accumulation and the efficiency with which ligand engagement is coupled to ZAP70 recruitment.

### Low-affinity CARs exhibit increased CD19-CAR cluster velocity compared to high-affinity CARs

We further quantified the velocity of CD19 clusters prior to ZAP70 colocalization, as well as the velocity of CD19-CAR/ZAP70 joint clusters during colocalization, using the CD19 signal as the reference (Figure 4a). The average CD19 velocity profiles from all trajectories of small clusters in maturing cells that started with no central cluster are shown in Figure 4b and Figure S10a for low- and high-CAR-expressing cells, respectively. As expected, CD19-CAR clusters moved consistently more slowly during ZAP70 colocalization than before ZAP70 recruitment, exhibiting a 0.7-0.8-fold reduction in speed (Figure S10b and Figure S11). This decrease in mobility likely reflects a combination of the increased effective size of the receptor complex upon ZAP70 recruitment and the additional molecular interactions associated with ZAP70 signaling. Binding of ZAP70 increases the molecular mass of the complex, which may reduce its mobility, while activation of bound ZAP70 results in its association and phosphorylation of substrates at the membrane, such as the key membrane-associated adaptor protein linker for activation of T cells (LAT) and SH2 domain-containing leukocyte protein of 76 kDa (SLP-76), which can be expected to reduce cluster diffusion speeds. When comparing CARs of different affinities, we found that, at high CAR expression levels, increasing CD19 density was associated with a slight increase in cluster velocity for both types of CARs Figure S10a,c), suggesting stronger activation is linked to increased cluster speeds. Interestingly, for low-CAR-expressing cells, increasing CD19 density was associated with faster movement of low-affinity CAT CAR clusters but slower movement of high-affinity FMC63 CAR clusters. As a result, the speed of low-affinity CAR clusters became up to 2-fold higher than that of high-affinity CAR clusters (Figure 4b,c. Thus, serial engagement of CD19 mediated by the higher off-rate of low-affinity CARs might also facilitate cluster reorganistion in the synapse by detachment/reattachment and thus ultimately result in faster CD19-CAR cluster movement.

**Figure 4.**
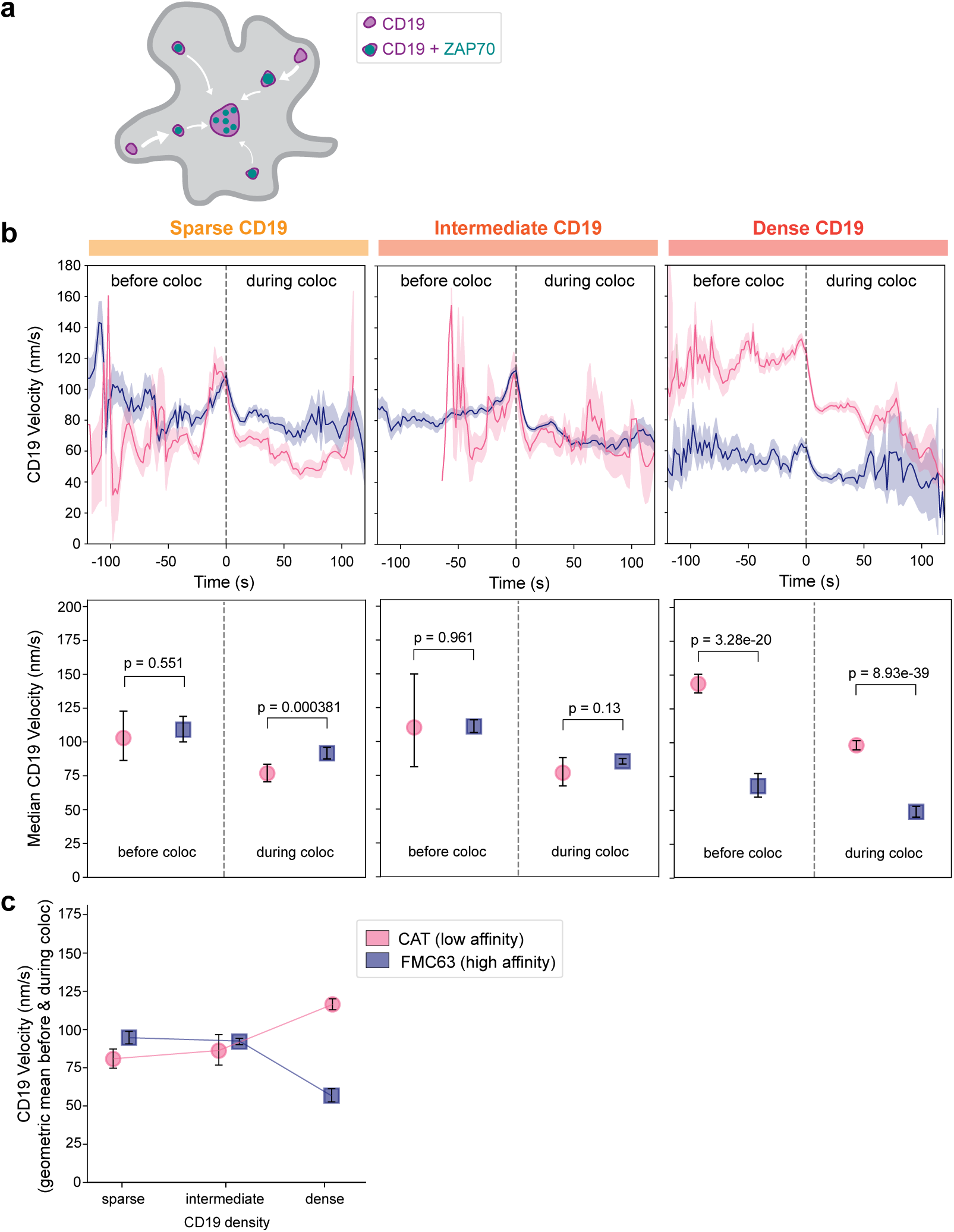
Low-affinity (CAT) CAR generated clusters moved 2-fold faster compared with high-affinity (FMC63) CAR-T cells expressing low level of CAR on dense CD19 SLBs. **(a)** Schematic of CD19-CAR microcluster motility before (CD19 only) and during ZAP70 colocalization (CD19 + ZAP70), thickness of the arrow represents the speed of the cluster. **(b)** CD19-JF646 velocity traces, shown as geometric means with SEM estimated from log-transformed values, from small CD19-CAR microcluster trajectories before and during colocalization with ZAP70, together with dot plots showing the corresponding median velocities with 95% nonparametric bootstrap confidence intervals, generated by low-affinity (CAT) and high-affinity (FMC63) CAR-T cells expressing low CAR levels interacting with sparse, intermediate, or dense CD19 SLBs. **(c)** Mean velocity of CD19-CAR microclusters across the full trajectory, including periods before and during ZAP70 colocalization. Values represent the geometric mean of the instantaneous CD19-JF646 microcluster velocities, with 95% nonparametric bootstrap confidence intervals. Number of analyzed tracks under different CD19 densities: sparse CD19, CAT (n = 55) and FMC63 (n = 218); intermediate CD19, CAT (n = 29) and FMC63 (n = 784); dense CD19, CAT (n = 447) and FMC63 (n = 147).

One of the downstream effects of TCR engagement is the initiation of actin cytoskeletal polymerization at the immunological synapse^25–28^. To determined whether CAR cell engagement with CD19 produces actin dynamics that drives movement of CAR microclusters, we visualized the spatiotemporal organization of F-actin together with ZAP70 at the immune synapses. We found that, upon CAR engagement with CD19, a peripheral actin-rich ring formed and subsequently moved inward along retrograde F-actin flow (Figure S12). ZAP70 microclusters formed in the peripheral ring, and both structures exhibited coordinated centripetal motion, consistent with previous observations in TCR signaling^27,29,30^.

Overall, these results suggest that at low expression levels, low-affinity CARs not only show more efficient CD19 binding and ZAP70 recruitment under dense CD19 conditions, but also display faster cluster motion than high-affinity CARs, coinciding with retrograde F-actin flow. These observations are likely mechanistically linked: ZAP70 recruitment and activation may increase cluster speed by activation of the actin cytoskeleton, while subsequent association with other signaling proteins reduces speed after the first peak, with increased ZAP70 recruitment for low-affinity CARs resulting in overall higher cluster speeds. Indeed at high expression levels, high affinity CARs also show higher velocity than the low expression cells due to their increased ZAP70 recruitment (Figure S8b,c and Figure S10c). However, they still display slower speeds than than the low-affinity CARs likely due to steric hindrance caused by a high amount of high-affinity receptor-ligand interactions in these clusters. Thus, our results show that CD19 affinity, along with CAR density and membrane expression, affects not only CD19 engagement and ZAP70 recruitment but also the dynamics of early signaling-cluster motion.

### Increased centripetal motion once ZAP70 colocalizes with CD19-CAR clusters

Next, we assessed how CAR affinity influences the inward movement of CAR signaling clusters by quantifying their directionality relative to the cell centroid. For each CD19-CAR microcluster trajectory, we defined a displacement vector from the start to the end of the trajectory and a radial vector from the trajectory start to the cell centroid (Figure S13a). Directionality was quantified as the cosine of the angle between these two vectors, yielding values from -1 to +1. Positive values indicate movement toward the cell center, with +1 representing purely centripetal motion; values near 0 indicate movement perpendicular to the radial direction; and negative values indicate outward movement, with -1 representing purely centrifugal motion.

In low-CAR-expressing cells with maturing synapses, directionality values were approximately 0.2-0.5 before ZAP70 colocalization for both CAR of different affinities, showing little directed movement to the synapse center (Figure S13b). However, during ZAP70 colocalization, these directionality increased to 0.5-0.7. A similar change in directionality was observed for high-CAR-expressing cells with mature synapses. In all cases, CD19-CAR clusters became more centripetal upon ZAP70 recruitment (Figure S13c). This demonstrates that, similar to TCR microcluster centripetal motion, the inward movement of CD19-CAR signaling clusters that drives synapse maturation requires ZAP70 recruitment. However, we observed little difference in cluster directionality for CARs of different affinities, likely because these analyses were performed on cells that formed mature synapses and hence pre-selected for cells with clusters showing centripetal motion.

When we focused our analysis on cells with disorganized synapses, we found that these exhibited decreased CD19-CAR cluster mobility (Figure S14a,b). Across all tested conditions for cells with disorganized synapses, low-affinity CARs moved consistently faster than high-affinity CARs, similar to what was observed for cells with mature synapses under the same conditions (Figure S10c), particularly at dense CD19. However, in low-CAR-expressing cells with disorganized synapses, ZAP70 colocalization did not typically enhance centripetal directionality, but cluster movement was directional, with direction values typically between 0.4 and 0.7, at all densities of CD19 (Figure S13d,e). Taken together, although most clusters moved more slowly in disorganized synapses, they still tended to move centripetally. This indicates that although ZAP70-driven centripetal motion can occur in disorganized synapses, the decreased mobility and the reduced recruitment of ZAP70 resulted in their failed ability to form mature synapses (Figure S14a,b, Figure 2c).

### Low-affinity CAR-T cells show more robust Ca^2+^ flux at dense CD19 conditions

Next, we tested if the increased ZAP70 recruitment efficiency and microcluster dynamics observed for low-affinity CAR-T cells also resulted in enhanced activation of downstream signaling pathways by quantifying the amount of calcium released upon CAR activation. Elevation of intracellular free calcium (Ca^2+^) is widely known to be the key requirement for successful T cell activation in response to antigen and other stimuli that engaged with the TCR^31^. Similarly, CD19 engagement by CAR is expected to initiate Ca^2+^ signaling that commits T cells to full activation and effector function^10^ (Figure 5a). To follow intracellular Ca^2+^ release upon synapse formation, we loaded cells with the calcium-sensitive dye Fluo-4 AM^32^. Depending on the strength of the Ca^2+^ flux and the cell’s ability to restore Ca^2+^ homeostasis, we observed different Ca^2+^ signaling responses, categorized by the duration for which intracellular Ca^2+^ levels remained elevated: sustained peaks gradually returned to baseline after reaching a maximum and remained elevated for an extended period, sharp peaks dropped rapidly to baseline following an initial increase, and no peak occurred when the Ca^2+^ signal remained below the detection threshold (Figure 5b,c).

**Figure 5.**
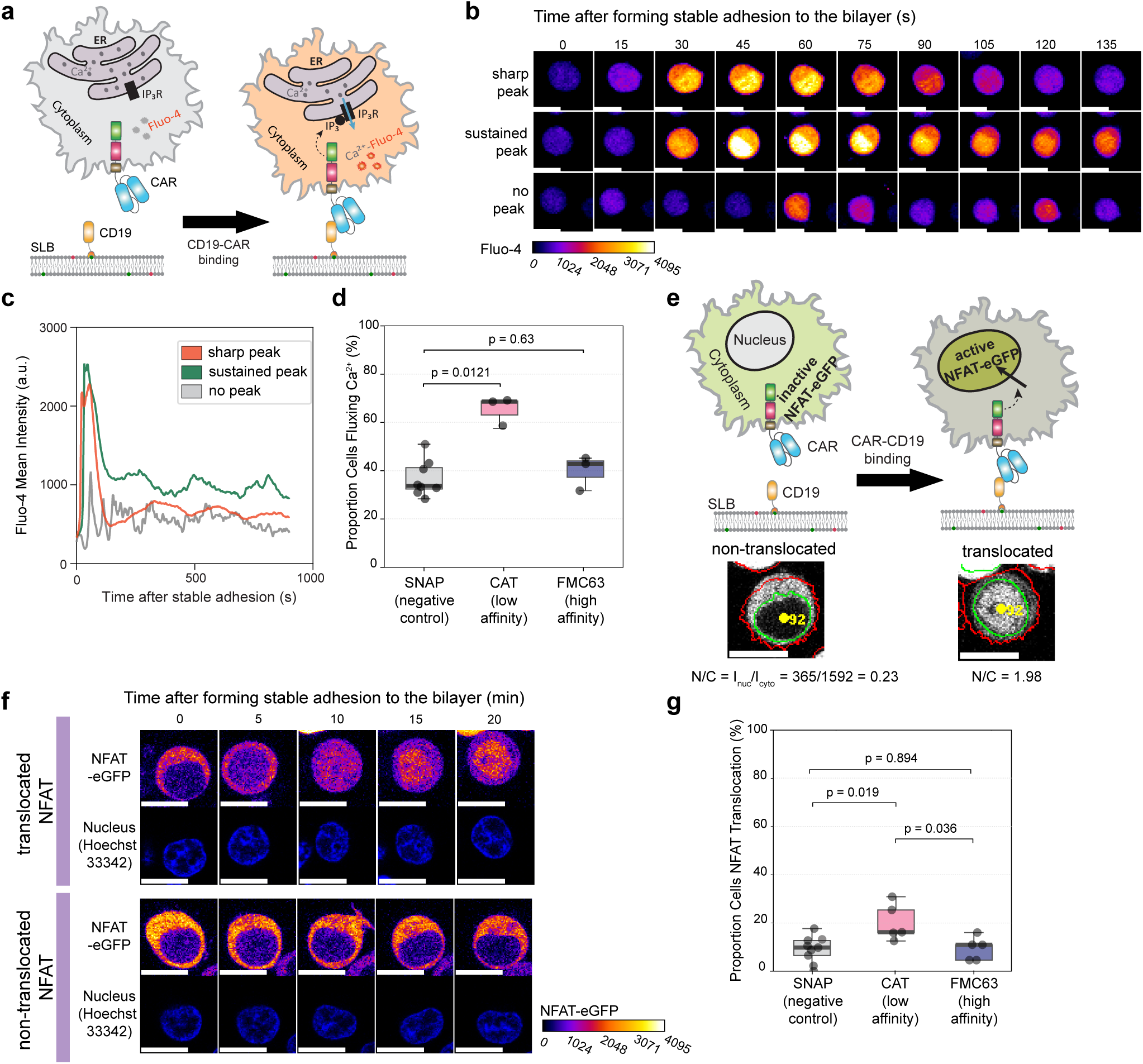
Enhanced Ca^2+^ flux and NFAT nuclear translocation in low-affinity (CAT) compared with high-affinity (FMC63) CAR-T cells at low CAR expression on dense CD19 SLBs. **(a)** Schematic illustrating intracellular Ca^2+^ signaling triggered by CD19-CAR engagement. CAR binding to CD19 on SLB induces IP_3_ production and activation of IP_3_ receptors on the endoplasmic reticulum (ER), resulting in Ca^2+^ release into the cytoplasm. Elevated cytosolic Ca^2+^ levels are detected as increased Fluo-4 fluorescence upon Ca^2+^ binding. **(b)** Representative time-lapse confocal images of single Fluo-4 AM-loaded cells, with Fluo-4 fluorescence serving as a readout of intracellular Ca^2+^ signaling. Cells exhibited distinct Ca^2+^ response profiles, characterized by a sharp peak (top), a sustained peak (middle), or no detectable (below-threshold) Ca^2+^ peak (bottom). Images were acquired during the first 135 s after stable cell contact with SLBs. Scale bars are 10 µm. **(c)** Corresponding mean Fluo-4 fluorescence intensity traces for the cells shown in (b), spanning the imaging period of 300 frames (900 s) used for analysis, including the interval beyond the initial 135 s displayed in (b). **(d)** Box plot showing the proportion (%) of CAR-T cells exhibiting robust intracellular Ca^2+^ flux upon interaction with CD19-presenting SLBs or with non-reactive SNAP-functionalized SLBs lacking CD19 as a negative control for low-affinity (CAT) and high-affinity (FMC63) CAR-T cells. Boxes indicate the median and interquartile range, and individual points represent independent replicates. Cell numbers: CAT (n = 375 cells, 3 replicates) and FMC63 (n = 308 cells, 3 replicates) on CD19 SLBs; SNAP control, combined CAT and FMC63 CAR-T cells (n = 654 cells, 8 replicates). **(e)** Schematic of NFAT-eGFP nuclear translocation induced by CD19-CAR engagement. Representative confocal images show a cell before CD19 engagement with NFAT-eGFP localized in the cytoplasm (non-translocated; N/C = 0.23) and after stable contact formation with the CD19 SLB, where NFAT-eGFP accumulated in the nucleus (translocated; N/C = 1.98). Nuclear boundaries (green; Hoechst 33342 staining) and whole-cell boundaries (red; NFAT-eGFP signal) were automatically segmented, and the cytoplasmic compartment was defined as the region between them. NFAT translocation was quantified using the nuclear-to-cytoplasmic mean fluorescence intensity ratio (N/C) and scored as positive when N/C *>* 1.1 for at least five consecutive frames. Scale bars are 10 µm. **(f)** Representative 20-min time-lapse confocal images showing a CAR-T cell with (top) or without (bottom) NFAT-eGFP nuclear translocation. Cell nuclei were visualized using Hoechst 33342 staining. Scale bars are 10 µm. **(g)** Box plot showing the proportion (%) of CAR-T cells exhibiting NFAT nuclear translocation upon interaction with CD19-presenting SLBs or with non-reactive SNAP-functionalized SLBs lacking CD19 as a negative control for low-affinity (CAT) and high-affinity (FMC63) CAR-T cells. Boxes indicate the median and interquartile range, and individual points represent independent replicates. Cell numbers: CAT (n = 70 cells, 5 replicates) and FMC63 (n = 25 cells, 5 replicates) on CD19 SLBs; SNAP control, combined CAT and FMC63 CAR-T cells (n = 43 cells, 9 replicates).

Across all conditions, both low- and high-affinity CAR-T cells interacting with CD19-SLBs or non-reactive SNAP-functionalized SLBs lacking CD19 (negative control) exhibited all three Ca^2+^ signaling patterns, albeit in different proportions. By quantifying Ca^2+^ signaling across hundreds of cells per condition, we determined the proportion of cells exhibiting significant Ca^2+^ flux, defined as either sustained or sharp peaks exceeding 35% above its lower baseline. In low-CAR-expressing cells interacting with dense CD19, 65% of cells expressing low-affinity CARs exhibited robust Ca^2+^ flux, in contrast to 40% of high-affinity CAR cells and 37% in the negative control (Figure 5d), indicating more robust CAR signaling in low-affinity CAR-T cells under these conditions. At high CAR expression levels, cells displayed equally strong Ca^2+^ flux for low-affinity and high-affinity CARs (Figure S15a).

Additionally, cells in the negative-control showed delayed calcium release than low-CAR-expressing cells of any affinity interacting with dense CD19 SLBs (negative control: 140 s, low expression CAT CAR: 99 s, low expression FMC63 CAR: 85 s, high expression CAT CAR: 80 s, high expression FMC63 CAR: 78 s, Figure S15b). Moreover, only 68% of Ca^2+^ fluxing cells in the negative-control condition exhibited sustained elevation (Figure S15c). In contrast, CAR-T cells engaging CD19 showed substantially higher frequencies of sustained elevation, with 80% for low CAR expression and 90-91% for high CAR expression across both affinities. These results show that CD19-CAR engagement drives a faster and more sustained Ca^2+^ elevation than conditions where CAR did not bind CD19.

Taken together, these results indicate that, in line with the CD19-CAR/ZAP70 findings, low-affinity CARs at dense CD19 again displayed stronger early CAR-signaling activation, including their intracellular Ca^2+^ responses. Of note, high-CAR-expressing cells generally initiated Ca^2+^ flux slightly earlier and sustained it for slightly longer periods compared to low-CAR-expressing cells, in line with a higher number of receptors being activated under those conditions.

### Low-affinity CAR-T cells show higher transcriptional activation than high-affinity CARs at dense CD19 levels

Translocation of NFAT to the nucleus, particularly NFATc2, is a key Ca^2+^-dependent step that initiates transcription of immune-regulatory genes^33^. To assess if NFAT translocation followed the trend observed for Ca^2+^ release, we introduced an NFAT-eGFP reporter into our CAR-T cells and monitored its cytoplasmic-to-nuclear translocation as a readout of CAR activation on CD19-functionalized SLBs (Figure 5e,f). Analysis of NFAT-eGFP translocation in low-CAR-expressing cells revealed that 20% of low-affinity CAR-T cells underwent robust NFAT nuclear translocation. In contrast, both the negative-control condition (CAR-T cells on non-reactive SNAP-SLBs) and high-affinity CAR-T cells engaging dense CD19 showed markedly lower frequencies of NFAT translocation (10% in both cases) (Figure 5g). In high-CAR-expressing cells, the fraction of cells displaying NFAT translocation was slightly increased to 27% and 20% for low- and high-affinity CAR, respectively (Figure S15d). This trend correlates well with our earlier observations showing the proportion of cells with strong Ca^2+^ release. Additionally, consistent with the median time to the first Ca^2+^ release (Figure S15b), the median time to NFAT translocation was markedly faster for any CAR-expressing condition compared to the negative control (Figure S15e) in high-CAR-expressing cells.

## Discussion

CAR affinity is frequently tuned to improve therapeutic performance, yet the mechanisms linking antigen binding to early signaling remain unclear. In this study, we find that at low expression levels, low-affinity CAT CARs form microclusters that recruit higher levels of ZAP70 while engaging fewer CD19 molecules than their higher-affinity counterpart. This indicates that phosphorylation of low-affinity receptors occurs more efficiently following ligand binding. Our observations support a model in which the low affinity interaction of CAT CARs allows for unbinding from CD19 more readily, analogous to TCR-pMHC interactions characterized by rapid dissociation kinetics. Such dynamics could facilitate serial triggering allowing a single CD19 molecule to sequentially engage multiple CARs and thereby amplify receptor activation and sustain ZAP70 recruitment within signaling microclusters (Figure 6a). Additionally, the lower number of receptor-CD19 binding interactions within the low-affinity CAR microclusters results in looser cluster organization, which may increase the accessibility for receptor phosphorylation by Src family kinases such as Lck, as indicated by the higher Zap70 binding. Conversely, with the exception of the high ligand density and receptor expression condition, the high-affinity interaction between FMC63 CAR and CD19 may constrain receptor-ligand turnover, limiting serial triggering and resulting in reduced ZAP70 recruitment relative to the level of CD19 engaged (Figure 6b)

**Figure 6.**
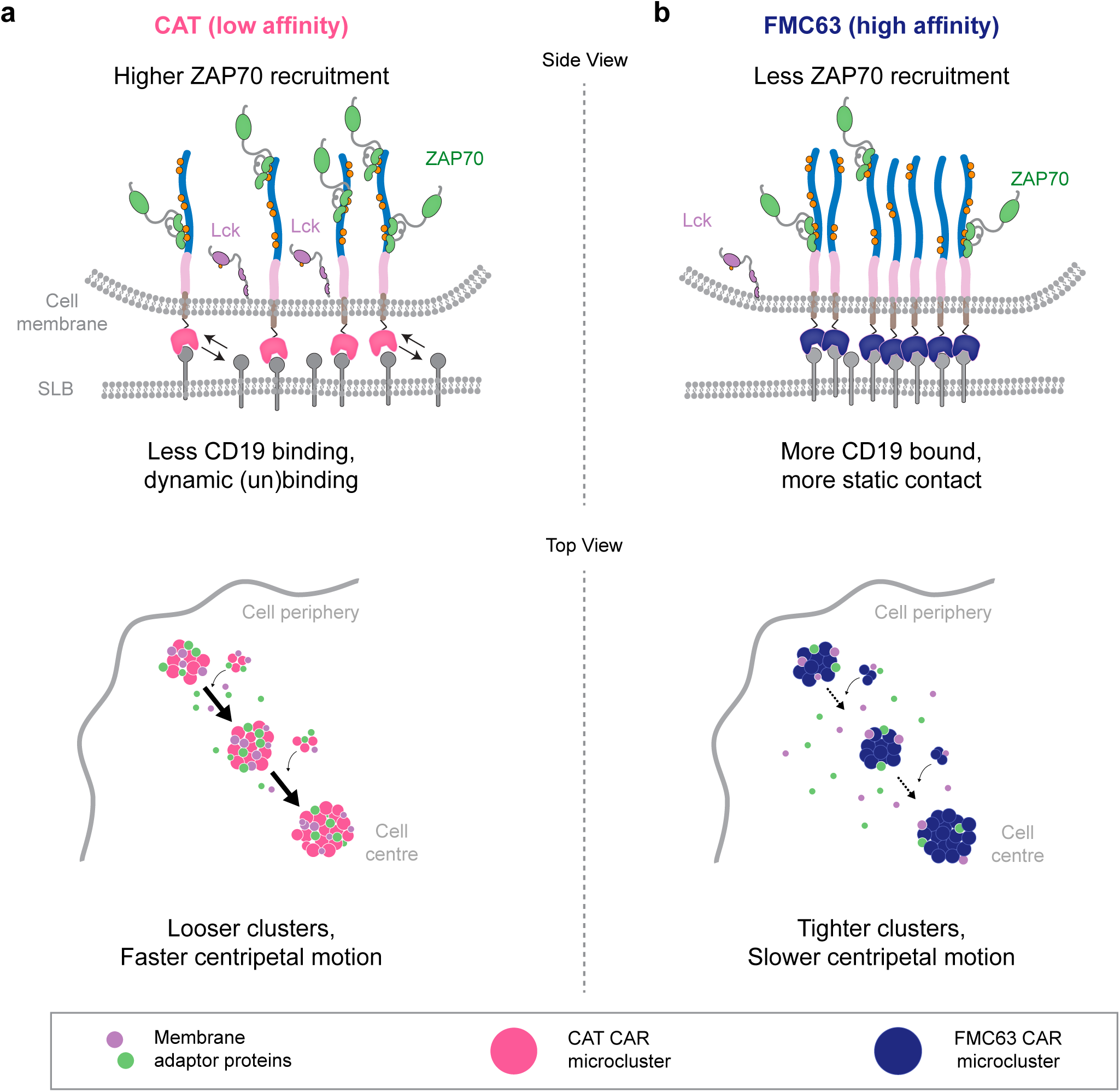
Schematic model showing how receptor kinetics, microcluster organization and motility enhance signaling and therapeutic efficacy of low-affinity CAT CARs. **(a)** Low-affinity CAT CARs form loose, highly motile microclusters that readily dissociate from CD19, potentially enabling serial triggering, sustained ZAP70 recruitment and enhanced integration of signaling from transmembrane adaptor proteins such as LAT and accessibility for phosphorylation by kinase Lck. **(b)** In contrast, high-affinity FMC63 CARs form tighter CD19 interactions that may restrict receptor turnover and microcluster motility, limiting serial triggering, adaptor protein recruitment, and sustained receptor phosphorylation, thereby reducing downstream T cell signaling.

Consistent with this model, low-affinity CAT CAR microclusters exhibited greater motility than high-affinity FMC63 CAR microclusters, moving approximately 2-fold faster in low-CAR-expressing CAR-T cells interacting with dense CD19 SLBs (54-113 molecules/µm^2^). This suggests that stronger interactions with CD19 and increased receptor binding result in more restricted centripetal movement. Additionally, low-affinity CAT CARs demonstrated a higher fraction of mature synapse formation and a superior ability to trigger downstream intracellular Ca^2+^ release and NFAT nucleus translocation. Taken together, these results demonstrate that low-affinity CAT CARs outperformed high affinity FMC63 CAR-T cells with the differential antigen affinities having consequences on receptor activation, immunological synapse formation, actin-driven centripetal movement and downstream cellular processes. We speculate that the increased motility of low-affinity CARs, together with their loose microcluster organization, enhances their ability to associate with, and integrate signaling from crucial adaptor proteins on the membrane, such as LAT, over time and across the synaptic interface. This would promote the observed efficient propagation of downstream T cell signaling responses despite lower levels of ligand engagement^34–36^. The stronger Ca^2+^ and NFAT responses in low-affinity CAT CAR-T cells may help explain their enhanced proliferation, cytotoxicity, expansion and antitumor activity reported previously, particularly against targets with high CD19 expression similar to the dense condition used in our experiments^4,34^. Conversely, high-affinity FMC63 CARs generate tighter, more static clusters that bind more CD19 yet recruit less ZAP70 and undergo slower centripetal motion. These properties may reduce association with membrane adaptor proteins, limiting downstream signaling pathway activation and prolonging exposure to the phosphatase CD45, a negative regulator of TCR-CD3*ζ* signaling^37^, at the periphery of the synapse, thereby limiting sustained receptor phosphorylation and ZAP70 recruitment.

The observed synapse phenotype was strongly shaped by antigen and receptor abundance. Increasing either CD19 density or CAR expression promoted mature over disorganized synapses formation, consistent with the number of receptor-ligand interaction influencing synapse organization. The clearest advantage of low-affinity CAT CARs emerged at dense CD19 and low CAR expression. At sparse CD19 densities (0.77-1.35 molecules/µm^2^), both CARs showed detectable antigen binding and ZAP70 recruitment, and differences in the recruitment efficiency and cluster velocity were small, with some measures slightly favoring high-affinity FMC63 CARs. This agrees with previous observations that CAT and FMC63 CARs display comparable killing of low-CD19 targets^4^. High CAR expression reduced the affinity-dependent difference in Ca^2+^ signaling, indication the the benefit of lower affinity CARs appears greatest when antigen is abundant relative to receptor availability.

Overall, our results support a model where low-affinity CAT CARs have enhanced signaling capacity due to microcluster organization and motion, which may allow for greater association with signaling adaptors and sustained receptor phosphorylation. Thus, weaker CAR-ligand interactions paradoxically promote more productive T cell activation than higher affinity binding, which could explain their improved therapeutic performance.

## Acknowledgements

We acknowledge Christian Niederauer for the initial development of single molecule tracking and calcium flux analysis pipelines, and for insightful discussions on microscopy and experiments. We thank Elisa Teunissen for assistance with viral transduction, Vanda Sunderlíková for providing the NFAT-eGFP construct, and Laura Weimann for providing the MATLAB script used for quantification of labeling stoichiometry for photobleaching step analysis. M.V.F. acknowledges support from ZonMw through an Off Road grant (project no. 04510252520069). This publication is part of project number VI.Vidi.203.037 of the Talent Programme which is financed by the Dutch Research Council NWO (C.N, K.A.G).

## Author contributions statement

Chikim Nguyen designed and performed the experiments, analyzed the data, and wrote the original draft and final versions of the manuscript. Gerard Castro-Linares developed the analysis pipelines, analyzed the data, and wrote analysis methods and revised the manuscript. Lucrezia Gatti assisted with cell sorting and flow cytometry data analysis. Zsolt Sebestyen provided expertise and support in flow cytometry. Ricardo A. Fernandes provided the CAR constructs. Sander J. Tans contributed scientific input throughout the project and revised the manuscript. Megan Farrell contributed to shaping the project, developing the model, and revising the manuscript. Kristina A. Ganzinger conceived and supervised the study, secured funding, provided scientific guidance, contributed to the original draft and revisions of the manuscript. All authors reviewed, revised, and approved the final version of the manuscript.

## Methods

### Constructs for lentiviral gene transfer

CAR plasmid constructs were synthesized by Integrated DNA Technologies (IDT) and cloned into the pHR lentiviral backbone (Addgene plasmid #79121). Correct assembly was confirmed by MluI/NotI restriction enzyme digestion and Sanger sequencing. Lentiviral particles were produced using the packaging plasmids psPax2 (Addgene plasmid #12260) and pMD2.G (Addgene plasmid #12259). Cell lines used in this study included Jurkat T cells (E6-1; ATCC TIB-152), HEK293T cells (ATCC CRL-3216), and Lenti-X 293T cells (Takara Bio, #632180).

### Generation of CAR-T cells

Jurkat cells were transduced with lentivirus produced in HEK293T or Lenti-X 293T cells. Packaging cells were seeded at 0.6 10^6^ cells per well in 6-well plates and transfected the following day at 60-80% confluency with a pHR transfer plasmid encoding the protein of interest (e.g., CAR, ZAP70-eGFP, or NFAT-eGFP; 0.75 µg), psPax2 (0.25 µg; 12260, Addgene), and pMD2.G (0.26 µg; 12259, Addgene) using GeneJuice (70967, Sigma-Aldrich) or FuGene 4K (E5911, Promega). After 72 h, viral supernatants were harvested, clarified by centrifugation and filtration, and incubated with 1 × 10^6^ Jurkat cells for 24 h. Transduced cells were subsequently washed and cultured in fresh medium. Four days post-transduction, Trypsin-EDTA (392-0435, VWR) was added directly to the cells at a 1:10 dilution (v/v) and incubated for 15 min. The cells were subsequently washed, and cultured in fresh medium for an additional 3 days. Cells were used for experiments or cryopreserved 7 days after transduction.

### Fluorescence-Activated Cell Sorting (FACS) for CAR expression

Transduced CAR Jurkat T cells were prepared for sorting by incubation with the human CD19 CAR Detection Reagent conjugated to biotin (130-129-550, Miltenyi Biotec) according to the manufacturer’s instructions. Briefly, 1 × 10^6^ CAR T cells in suspension were centrifuged at 300 x g for 5 min, and the resulting cell pellet was resuspended in PBS buffer containing 10% FBS (A3160802, Gibco). The human CD19 CAR Detection Reagent was diluted 1:50 from stock and incubated with the cells for 10 min in the dark at room temperature. Subsequently, cells were washed twice with 1 mL PBS/10% FBS buffer by centrifugation at 300 x g for 5 min.

A second staining step was performed by incubating cells with an antibody mixture containing Streptavidin-PE conjugate antibody (36777, Cell Signaling) and pan*αβ* TCR-APC (IP26, eBioscience) for 30 min at 4 *^◦^*C. Cell sorting was carried out using the BD FACSAria II Cell Sorter. The FACS gating strategy was applied sequentially: first, live cells were identified based on size and granularity using FSC/SSC parameters; second, single cells (singlets) were selected through FSC discrimination; third, populations with comparable *αβ* TCR expression were gated; and finally, cells were segregated into high- and low-CAR expression subsets according to CD19-PE mean fluorescence intensity (MFI). Transduced cells were selected based on comparable pan*αβ* TCR-APC signal (MFI of 1-5 × 10^3^) and subsequently divided into two populations according to CD19-PE fluorescence, indicative of CAR expression at the cell membrane: high CAR expression (CD19-PE high; MFI *>* 1-2 × 10^4^) and low CAR expression (CD19-PE low; MFI *<* 1 × 10^3^). The two sorted populations were collected in 5 mL FACS tubes containing 500 µL RPMI-1640 medium supplemented with 300 U/mL penicillin/streptomycin and 10% FBS.

Sorted cells were cultured for the first 2 days at a concentration of 0.5 × 10^6^ cells/mL in RPMI-1640 medium containing 300 U/mL penicillin/streptomycin and 10% FBS. on the second day after the sorting, 50 IU/mL of interleukin-2 (IL-2, Clinigen) was added to the cell culture to support survival and modest proliferation. Cells were passed at least 3 times before being used for experiment.

### Protein labeling

Janelia Fluor 646 (JF646; 6148, Tocris) NHS ester was dissolved in anhydrous DMSO to 10 mM. Human CD19 containing a 10 × His tag (500 µg; CD9-H52H2, ACROBiosystems) was reconstituted in 250 µL sodium borate buffer (50 mM, pH 8.5; 1330-43-4, Thermo Scientific Chemicals) and incubated for 1 h at room temperature. JF646 (23.43 µL, 10 mM; 15:1 dye-to-protein molar ratio) was added, and the reaction was incubated for 1 h at room temperature in the dark. Excess dye was removed using a 7kDa MWCO Zeba spin desalting column (89890, Thermo Scientific) according to the manufacturer’s instructions. The purified CD19-JF646 was snap-frozen in liquid nitrogen and stored at -80 *^◦^*C until use.

The labeling stoichiometry of JF646-labeled CD19 was quantified by single-molecule photobleaching step counting. Proteins were immobilized on piranha-cleaned and plasma-cleaned coverslips at low surface densities (0.1-0.2 molecules*/*µm^2^) and imaged by TIRF microscopy under high-intensity 638 nm excitation until *>*90% of fluorophores were photobleached. Images were acquired with 50-500 ms exposures over 1000-1500 frames. Labeling stoichiometry was determined using a custom MATLAB script (available upon request) for photobleaching step analysis.

### SLB preparation

Lipids dissolved in chloroform were mixed at a molar ratio of 97.5% DOPC (850375, Avanti), 1% DGS-NTA(Ni) (790404, Avanti), 1% MCC-DOPE (780201, Avanti), and 0.5% DOPE-ATTO390 (390-161, ATTO-TEC). Chloroform was removed under a nitrogen stream, and lipid films were vacuum-desiccated overnight. The dried films were hydrated in HEPES-buffered saline (HBS; 20 mM HEPES (15630080, Thermo Fisher Scientific), 140 mM NaCl (310166, Sigma-Aldrich), pH 7.4-7.6) to a final lipid concentration of 2 mg/mL and vortexed for 5 min to generate multilamellar vesicles (MLVs), which were aliquoted and stored at -20 *^◦^*C until further use. For preparation of small unilamellar vesicles (SUVs), MLVs were thawed and sonicated on ice for 15 min until the suspension became transparent. SUVs were subsequently diluted to (0.2 mg/mL) and used on the same day.

Piranha-cleaned glass coverslips (22 × 22 mm, 1.5H) were assembled into chambers, plasma-cleaned for 10 min, and incubated with SUVs (0.2 mg/mL, 30 min) to allow spontaneous SLB formation. Bilayers were washed with HBS and blocked for non-specific binding with 0.5% (w/v) casein (C3400, Sigma-Aldrich) in HBS supplemented with 20 mM MgCl_2_ for 30 min. 10xHis-tagged ICAM-1 (12.8 nM), 10xHis-tagged CD19 (variable concentrations), or 10xHis-tagged SNAP protein (48.5 nM) were then added and allowed to bind DGS-NTA(Ni) lipids for 1 h. After two washing steps with the same blocking solution, SLBs were stored in a dark, humidified chamber and imaged on the same day.

### TIRF microscopy

Fluorescence imaging was performed on a custom-built TIRF microscope, described before by Niederauer et al.^38^. Briefly, the microscope was equipped with a 60× oil-immersion objective (NA 1.49, Nikon). Excitation was provided by a multi-line laser combiner delivering 405 nm, 488 nm, 561 nm, and 638 nm laser lines. The excitation beam was fiber-coupled, spectrally cleaned, beam-shaped, and focused onto the back focal plane of the objective, allowing adjustment between widefield, HILO, and TIRF illumination by lateral displacement of the beam. Fluorescence emission was separated using multi-band dichroic mirrors and bandpass filters and relayed through a 4 *f* imaging system onto distinct regions of a single sCMOS camera (Prime BSI, Teledyne Photometrics) for simultaneous multi-color detection. Microscope control and data acquisition were performed using custom software.

CAR-T cells expressing ZAP70-eGFP were generated by lentiviral transduction and sorted to obtain populations with comparable *αβ* TCR levels and either low or high CAR expression. For each imaging chamber, 300,000 cells were incubated with 0.3 µM Chol-DNA for additional 10 min at room temperature and then washed twice with imaging buffer (phenol-red-free RPMI 1640 supplemented with 1% (v/v) sodium pyruvate, 1% (v/v) HEPES, and 1% (v/v) GlutaMAX). Cells were resuspended in 100 µL imaging buffer and introduced into chambers containing functionalized SLBs. Imaging was performed at 37 °C immediately after cell addition. Images were acquired using TIRF microscopy, with objective heating set to 40 °C; after heat dissipation, the temperature at the glass-sample interface stabilized at approximately 37 °C. ZAP70-eGFP and CD19-JF646 were excited using 488 nm and 638 nm laser, respectively, with a 500 ms exposure time and a 2 s frame interval.

SLB fluidity was checked by fluorescence recovery after photobleaching (FRAP) of DOPE-ATTO390 lipids within the SLBs using 405 nm laser.

Chol-DNA sequence (5*^′^* to 3*^′^* end): ACT GAC TGA CTG ACT GAC TG T TTT TTT TTT TTT TTT TTT TTT TTT TTT

TTT TTT TTT TTT TTT TTT TTT TTT TTT TTT TTT TTT TTT TTT TTT TTT TTTT-CholTEG

DNA-thiol sequence (5*^′^* to 3*^′^* end): CAG TCA GTC AGT CAG TCA GT-thiol C6

### Measurements of antigen densities

CD19-JF646 antigen densities on SLBs were quantified by either diffraction-limited spot counting or intensity-based analysis, depending on surface density. At low densities (*<*0.1-0.2 molecules*/*µm^2^), individual fluorophores were resolved and counted using Picasso Localize^39^, which detects single-molecule emission events using a gradient-based algorithm followed by least-squares Gaussian fitting. Molecule counts from the first five frames were averaged across ten regions of interest (ROIs) and normalized to ROI area (4357 µm^2^) to obtain fluorophore density.

At higher densities where individual molecules cannot be resolved easily, antigen density was estimated from fluorescence intensity following established methods with some adaptations^10,40,41^. Single-molecule intensities were determined from low-density SLBs using TrackMate (Fiji) with Laplacian-of-Gaussian detection, which reports raw pixel intensities. Integrated single-molecule intensity was calculated from around 2300 detected spots after background subtraction. For each ROI, integrated fluorescence intensity was obtained from the background-subtracted mean intensity multiplied by ROI area and divided by the single-molecule intensity to estimate molecule number. Fluorophore density was calculated by normalizing molecule counts to ROI area, and antigen density was obtained by correcting for the experimentally determined CD19 labeling stoichiometry determined above. Six to ten ROIs per SLB were analyzed, and their mean density was reported as the representative CD19 density.

### Image processing and single-cell ROI selection

Because multi-color emission was projected onto distinct regions of a single sCMOS sensor, the raw images were first spatially cropped to isolate the individual spectral channels. To ensure accurate channel alignment for colocalization analysis, we corrected for chromatic aberrations using affine transformation. Channel-to-channel registration was determined using TetraSpeck **(T7279, Thermo Fisher Scientific)** fluorescent bead images and the Descriptor-based registration (2D/3D) plugin^42^ in Fiji^43^, yielding affine transformation matrices for the left-to-center and right-to-center channel mappings. These transformation matrices were subsequently applied to the cropped raw images using the affine_transform function from scipy.ndimage.

Using the affine-transformed images, single-cell regions of interest (ROIs) were manually defined. ROIs were drawn around individual cells on maximum-intensity projections using the freehand selection tool in Fiji^43^. The resulting ROI masks were saved and used through subsequent analysis steps to uniquely identify and process individual cells within the same field of view.

### Synapse maturation analysis

To obtain an objective and quantitative measure of immune synapse maturation, we implemented a convolutional neural network-based classifier trained on TIRF microscopy images of ZAP70 recruitment. Individual synapse frames were manually annotated according to established morphological criteria reflecting the spatial organization and clustering of ZAP70, yielding 394 immature and 377 mature images. Frames were sampled from the full dataset of tracked cells and filtered to retain only time points separated by at least 30 frames within each cell, minimizing temporal redundancy. Classification criteria included the presence of large centralized clusters, their relative intensity, and the degree of clustering, distinguishing small dynamic microclusters from large stable central clusters^13^. Cropped single-cell regions of interest (ROIs) were resized to 224 × 224 pixels and intensity-normalized to the range 0-1. Because the model architecture was pretrained on RGB images, grayscale inputs were replicated across three channels. The dataset was divided into training (90%) and validation (10%) subsets using stratified sampling to preserve class balance.

Feature extraction was performed using a MobileNetV2 backbone pretrained on ImageNet^44^, followed by a task-specific classification head consisting of global average pooling, dropout (0.5), and a fully connected layer (64 units, ReLU activation) with a sigmoid output for binary classification. Training was performed using binary cross-entropy loss and the Adam optimizer in two stages: first with the convolutional backbone frozen for up to 20 epochs, and subsequently with the final five convolutional layers unfrozen for up to 30 additional epochs to allow limited fine-tuning of higher-level spatial features associated with synapse architecture. Early stopping and adaptive learning rate reduction based on validation loss were applied to prevent overfitting. Mild class imbalance was addressed using class weighting (1 for immature and 1.1 for mature). To enhance robustness to variability in cluster geometry and cell orientation, online data augmentation was applied to the training set, including random flips, affine shifts, zoom, shear, and rotations (±20°), with rotation-induced empty regions filled using noise sampled from the image intensity distribution to avoid artificial edge artifacts. Model performance was evaluated on the held-out validation set using accuracy, precision, recall, and area under the receiver operating characteristic curve (AUC). The trained classifier was subsequently used to quantify synapse maturation across experimental conditions. Source code and analysis scripts are available in GitHub^45^.

The classifier output for each frame was a probability *p_mature_*(*t*) ∈ [0, 1]. For each cell, the temporal trajectory *p*(*t*) was fitted with a sigmoidal function *p*^(*t*); when the fit achieved *R*^2^ *>* 0.75, classification was based on *p*^(*t*), otherwise on *p*(*t*) (smoothed with a moving average of 5 frames). The maturation transition time was defined as the first frame where *p*^(*t*) > 0.5. The initial and final probabilities, *p_start_* and *p_end_*, were defined as the mean probabilities over the first and last 5 frames of the selected trajectory, respectively, and *p_max_*and *p_min_* denote its maximum and minimum values. Cells were categorized according to threshold criteria on these quantities and the dynamic change Δ*p* = *p_end_ p_start_*, as summarized in Table 1. This classification was done using the predict_maturation function within the Cell_Analyzer class in the GitHub repository^45^. Non-classifiable synapses were removed from further analysis.

**Table 1.** Criteria for cell-level immune synapse maturation classification based on the maturation probability trajectory.

| Category | Classification Criteria |
| --- | --- |
| Never mature (disorganized) | $p_{\max} < 0.4$ |
| Matures (maturing: started without central cluster) | $p_{\text{start}} < 0.4$, $p_{\text{end}} > 0.6$, and $\Delta p > 0.4$ |
| Starts mature (maturing: started with central cluster) | $p_{\text{start}} > 0.6$ and $p_{\min} > 0.4$ |
| Not classifiable | All other cases |

### Large ZAP70 cluster analysis

Large ZAP70 clusters were detected and quantified using a custom python segmentation pipline^45^. For each manually defined single-cell ROI, image stacks were cropped to the ROI bounding box. Cluster segmentation was performed by combining a global and a local threshold. A global Li threshold was computed on the maximum-intensity projection of the stack and applied to each frame, producing a global binary mask after morphological closing. In parallel, an adaptive Phansalkar threshold was computed independently for each frame using a local window radius of 15 pixels with parameters p=2 and q=8. The final cluster mask was obtained as the logical AND intersection of the global and local masks (*M_clusters_* = *M_Li_^M_Phansalkar_*), followed by morphological closing/opening and removal of connected components smaller than 80 pixels in each frame.

Connected components were labeled in each frame and, for each cluster, several cluster morphological and intensity features were extracted, such as area, circularity, centroid position, integrated intensity, and median intensity, both not-normalized and normalized against the median intensity background outside of the cell ROI. In addition, per-cell, per-frame summary statistics were computed as weighted means of cluster features, with weights proportional to *area normalized median intensity*, to capture the dominant structural characteristics of the synapse at each time point.

### Small cluster (co-)localization and tracking

To trajectory diffraction-limited clusters of ZAP70 and CD19 in TIRF images, we used a custom-written Python pipeline^45^. Single-molecule localizations were obtained using Picasso v0.8.0^46^ with a box size of 7 pixels and a minimum gradient threshold of 500 for both channels. Localizations outside the manually defined ROIs were excluded, and each remaining localization was assigned an ROI index to enable single-cell analysis. Localizations within clusters were also removed since they only indicate heterogeneity inside the large clusters and not active signaling CD19-CART-ZAP70 small diffraction-limited clusters moving towards the larger clusters.

Trajectory reconstruction was performed using trackpy v0.5.0 with a search range of 15 pixels, memory of 1 frame, adaptive_step = 0.95, and adaptive_stop = 2 pixels. Only trajectories longer than 5 frames were considered for further analysis. ZAP70 and CD19 trajectories were classified as colocalized when their per-frame Euclidean distance was *<*250 nm (∼2 px center-to-center) for at least 5 consecutive frames. If multiple candidate trajectory pairs satisfied these criteria, only the pair with the smallest mean distance during the overlapping interval was retained.

### Analysis of the velocity, intensity, and directionality of small clusters

For each colocalized CD19-ZAP70 trajectory, instantaneous particle velocities were computed as frame-to-frame displacements divided by the corresponding time interval (2 seconds). Fluorescence intensity at each time point was quantified as the mean intensity within a 3 × 3 pixel patch centered at the particle position in the raw image and normalized to the median intensity of the corresponding frame to account for background signal. Median instantaneous velocity and intensity were calculated over the full trajectory or over defined temporal segments (before and during colocalization), depending on the analysis.

Directionality was quantified over defined temporal segments (before and during colocalization) by comparing the net displacement vector of each segment to the inward radial vector pointing toward the cell centroid (based on the ROI). For each segment (before and during colocalization) the displacement vector Δ**x** = **x**_2_ - **x**_1_ was compared to the inward radial vector **r** = **c** - **x**_1_, where **c** denotes the cell centroid. Directionality was defined as the cosine of the angle between these vectors, cos *θ* = Δ**x·r***/*( ||Δ**x||** ||**r**||), i.e., the dot product of the displacement and inward radial vectors normalized by the product of their magnitudes. cos *θ* has values ranging from +1, which indicates perfect inward motion, and -1, which indicates outward motion.

Finally, for the analysis of dynamics around colocalization events, CD19-ZAP70 co-trajectories were temporally aligned to the first frame of colocalization, defined as *t* = 0. Only colocalized trajectories lasting at least 5 frames and containing at least 2 frames preceding colocalization onset were included. Velocity and intensity values were retained only within the temporal window spanning the before colocalization period and the during colocalization interval itself. Instantaneous velocity and normalized intensity traces were then averaged using the geometric mean at each aligned time point. Variability was estimated from the standard error of the log-transformed values.

### Statistical analysis

The percentage (%) or the proportion of cells in each synapse maturation category was determined for each CAR-T cell subgroup (low or high CAR expression and low- or high-affinity CARs) under each CD19-density condition. Uncertainty was shown as the standard error of a binomial proportion, 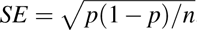, where *p* is the proportion of cells in a given category and *n* is the number of classified cells.

Velocity measurements were strictly positive and were therefore summarized using geometric means with 95% confidence intervals calculated on log-transformed values. Comparisons between FMC63 and CAT were performed using Welch’s two-sample t-test on log-transformed velocities. For paired before versus during colocalization velocity comparisons, fold changes were computed per co-trajectory as during/pre, log-transformed, and tested against no change using a one-sample t-test against zero; FMC63 and CAT fold changes were compared using Welch’s t-test on log fold changes.

Since directionality values were bounded between -1 and 1, they were summarized using the median with 95% non-parametric bootstrap confidence intervals based on 5000 resamples. To test whether directionality changed upon colocalization, the per-trajectory difference Δ*D* = *D_during_ D_before_* was calculated, and a one-sided paired sign-flip permutation test with 10,000 permutations was used to test whether the median Δ*D* was greater than zero.

Whole-synapse integrated intensities were also analyzed on a log scale and summarized as geometric means with 95% confidence intervals. Mature and non-mature synapses were compared within each CAR using Welch’s t-test on log-transformed integrated intensity values.

Small-cluster intensities were summarized using the median with 95% non-parametric bootstrap confidence intervals, because these distributions were not assumed to be normally distributed. FMC63 and CAT were compared within each timing window using a two-sided label-permutation test on the median difference with 10,000 permutations. For paired intensity fold changes between before and during colocalization, log fold changes were calculated per co-trajectory; within CAR-T changes were tested using a two-sided sign-flip permutation test, and FMC63 versus CAT differences were tested using a two-sided label-permutation test.

### Calcium (Ca^2+^) imaging

For each imaging chamber, 600,000 CAR-T cells were loaded with 2 µM Fluo-4 AM (6255, Tocris) for 30 min at 37 °C to visualize the spatial and temporal dynamics of cytoplasmic Ca^2+^ signaling. Half of the cells were subsequently incubated with 0.3 µM Chol-DNA for 10 min at room temperature and washed twice with imaging buffer. All cells were then resuspended in 100 µL imaging buffer and introduced into chambers containing functionalized SLBs. Confocal imaging (Nikon A1R-MP) was performed at 37 °C immediately after cell addition using 488 nm excitation and brightfield illumination with a 3 s frame interval.

### Ca^2+^ flux analysis

Candidate cells were detected in the first 300 frames (15 min) using a Difference-of-Gaussians (DoG) approach tuned to an expected cell diameter of approximately 12 *µ*m (allowing for 20% size variation and using a detection threshold of 3). For each detected blob at position (*x, y*) and scale *σ* , a circular region of interest with radius *r* =√2σ was used to extract the mean intensity, and subsequently tracked using trackpy with a search_range of 8px and a memory of 2 frames. Cell trajectories shorter than 75 frames (3.75 min or 225 s) were discarded. The mean intensity trace was then normalized to its maximum value. Peaks in the normalized mean intensity trace were identified using a scipy.signal.find_peaks, with constraints on minimum prominence (≥ 0.1), minimum height (≥ 0.2), and maximum width (≤ 50 *frames*). In scipy.signal.find_peaks, the peak prominence is defined as the vertical distance between the peak and the higher of its two surrounding baseline. This can underestimate the prominence of incomplete or asymmetric peaks, prominence was recalculated as the vertical distance between the peak and the median intensity of the lowest baseline region. The trace was then flagged as peak-containing when at least one peak was detected for that trajectory with a recalculated prominence higher than 0.35. We also extracted the time until the first peak as the time difference between the first detected peak and the start of the trajectory. Peaks were classified as complete when |(*R-L*)*/L| <* 1.0 (were *R* and *L* are the median intensities of the left and right baseline regions flanking the peak, respectively), indicating comparable baseline intensities on both sides of the peak. The code used for this analysis can be found in github^45^.

### NFAT imaging

For each imaging chamber, 300,000 CAR-T cells expressing NFAT-eGFP were incubated with 2 µM Hoechst 33342 (62249, Thermo Scientific) for 30 min at 37 °C to label nuclei for visualization and tracking. Cells were then incubated with 0.3 µM Chol-DNA for 10 min at room temperature, followed by two washes with imaging buffer. Cells were resuspended in 100 µL imaging buffer and introduced into chambers containing functionalized SLBs. Confocal imaging was performed at 37 °C, beginning 3 min after cell addition. Hoechst-stained nuclei and NFAT-eGFP were excited using 405 nm and 488 nm laser with a 30 s frame interval.

### NFAT analysis

Nuclei were segmented from the nuclear channel after illumination correction (division by a heavily blurred average projection, Gaussian *σ* = 100) and per-frame normalization. Segmentation was performed by Gaussian smoothing (*σ* = 1), triangle thresholding, morphological cleaning (removal of objects *<* 200 px, hole filling), and distance-transform watershed splitting (min_d_istance = 15 px). Cytoplasm regions were obtained from the fluorescence channel by triangle thresholding and morphological filtering, followed by assignment of pixels within a radius of 10 px to the nearest nucleus, yielding one cell region per nucleus. Nuclei were linked over time using trackpy (search_r_ange = 25 px, memory = 2), and trajectories shorter than one-third of the movie duration were discarded.

For each tracked cell, mean fluorescence intensities were extracted per frame in the nucleus and cytoplasm (excluding the nucleus) from the original image (not the illumination-corrected image), and the nuclear-to-cytoplasmic ratio was computed as N*/*C = *I*_nuc_*/I*_cyto_. A cell was classified as NFAT-translocating when N*/*C *>* 1.1 for at least 5 consecutive frames. The code used for this analysis can be found in github^45^.

---

**Figure S1.**
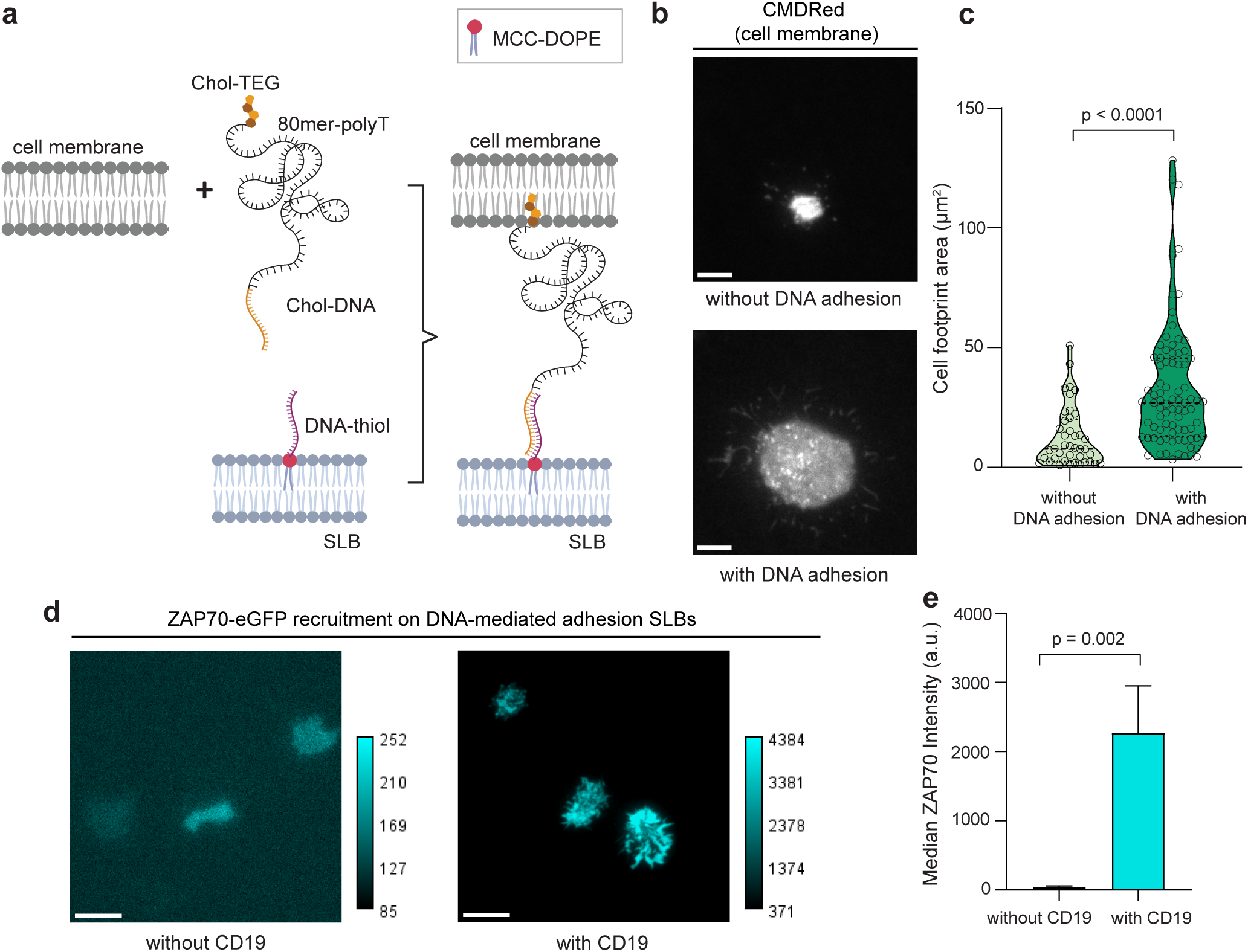
Synthetic DNA-mediated adhesion increased cell-SLB binding, producing approximately 2.5-fold larger contact footprints without non-sepcific ZAP70 recruitment. **(a)** Schematic of the synthetic DNA-mediated adhesion system. Cells were incubated with Chol-DNA, composed of a Chol-TEG moiety linked to an 80mer-polyT linker and a 20-nucleotide adhesive sequence (yellow). This sequence hybridizes with its complementary DNA-thiol strand (purple) pre-functionalized on the SLB, enabling specific synthetic adhesion between the cell membrane and the SLB. **(b)** Representative TIRF images of cell footprints labeled with CMDRed on DNA-thiol-functionalized SLBs in the presence and absence of the DNA-mediated adhesion system. Scale bars are 5 µm. **(c)** Violin plot showing the distribution of cell footprint areas for cells without and with DNA adhesion. Individual points represent individual cells. **(d)** Representative TIRF images (three cells per condition) of CAR-T cells on DNA-mediated adhesion SLBs with or without CD19. Scale bars are 5 µm. **(e)** Bar plot showing the median background-corrected ZAP70-eGFP fluorescence intensity at the CAR-T cell immune synapse in the absence or presence of CD19 on SLBs.

**Figure S2.**
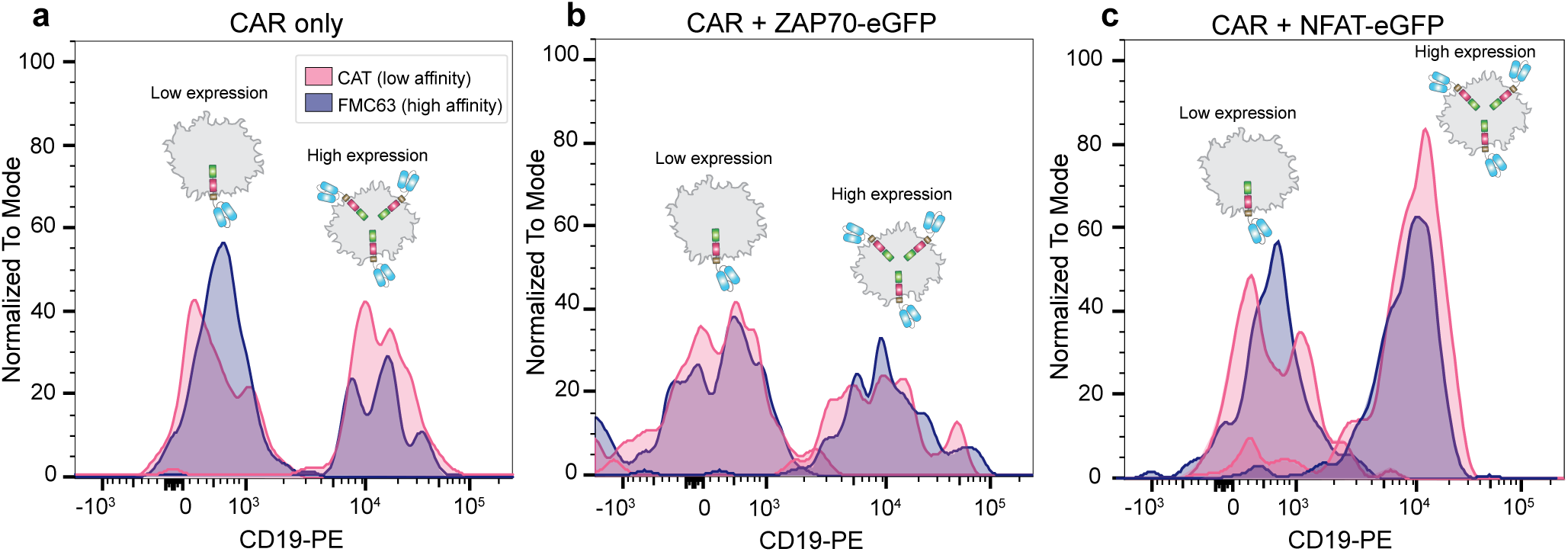
FACS analysis of CAR expression in CAR-T cell lines. CAR expression on the cell surface was quantified by CD19-PE staining in cells transduced with **(a)** CAR alone, **(b)** CAR and ZAP70-eGFP, or **(c)** CAR and NFAT-eGFP. Based on CD19-PE fluorescence intensity, cells were classified as low expression (CD19-PE MFI *<* 1×10^3^) or high expression (CD19-PE MFI *>* 1-2×-10^4^). The low- and high-CAR-expressing populations exhibited an approximately 10-20-fold difference in CD19-PE MFI, while maintaining comparable surface expression levels of *αβ* TCR.

**Figure S3.**
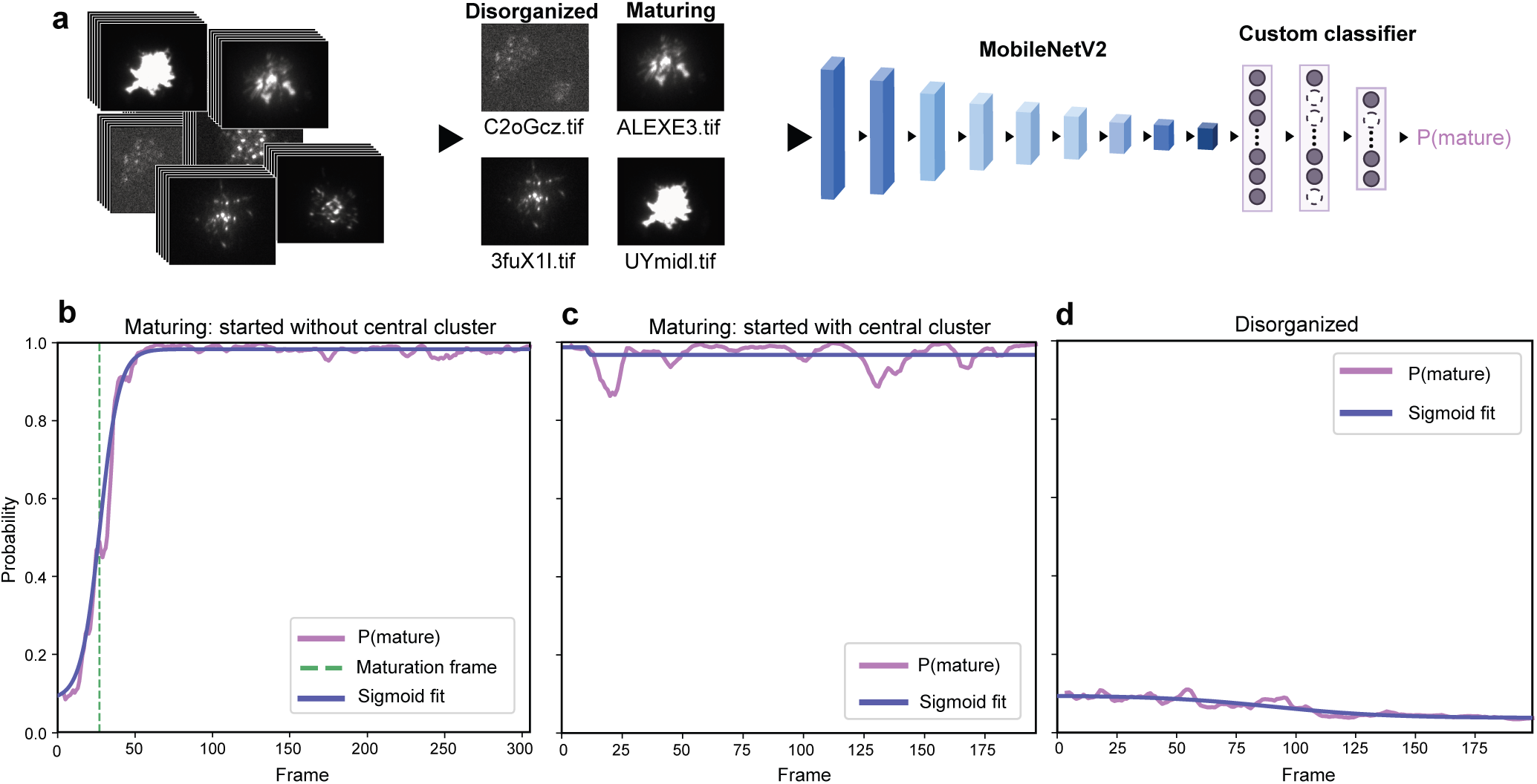
Classification pipeline for CAR immune synapse maturation based on ZAP70 recruitment dynamics. **(a)** From the raw TIRF videos of ZAP70 recruitment to CD19-CAR, 771 frames were randomly selected across different videos. When multiple frames were selected from the same video, they were separated by at least 30 frames, corresponding to 1 min. All selected frames were assigned randomized file names to anonymize them and minimize classification bias. These isolated frames were manually classified as either maturing synapses, 377 images, or disorganized synapses, 394 images, based on morphological criteria. A convolutional neural network based on MobileNetV2^44^ was then trained to output the probability that a single frame contained a maturing synapse. Additional details on the training parameters are provided in the Methods section. Once trained, the classifier was applied to full videos to obtain the probability of synapse maturation over time, as shown in panels (b-d). The resulting traces were classified as: **(b)** maturing synapse started without central cluster, **(c)** maturing synapse started with central cluster, and **(d)** disorganized synapse.

**Figure S4.**
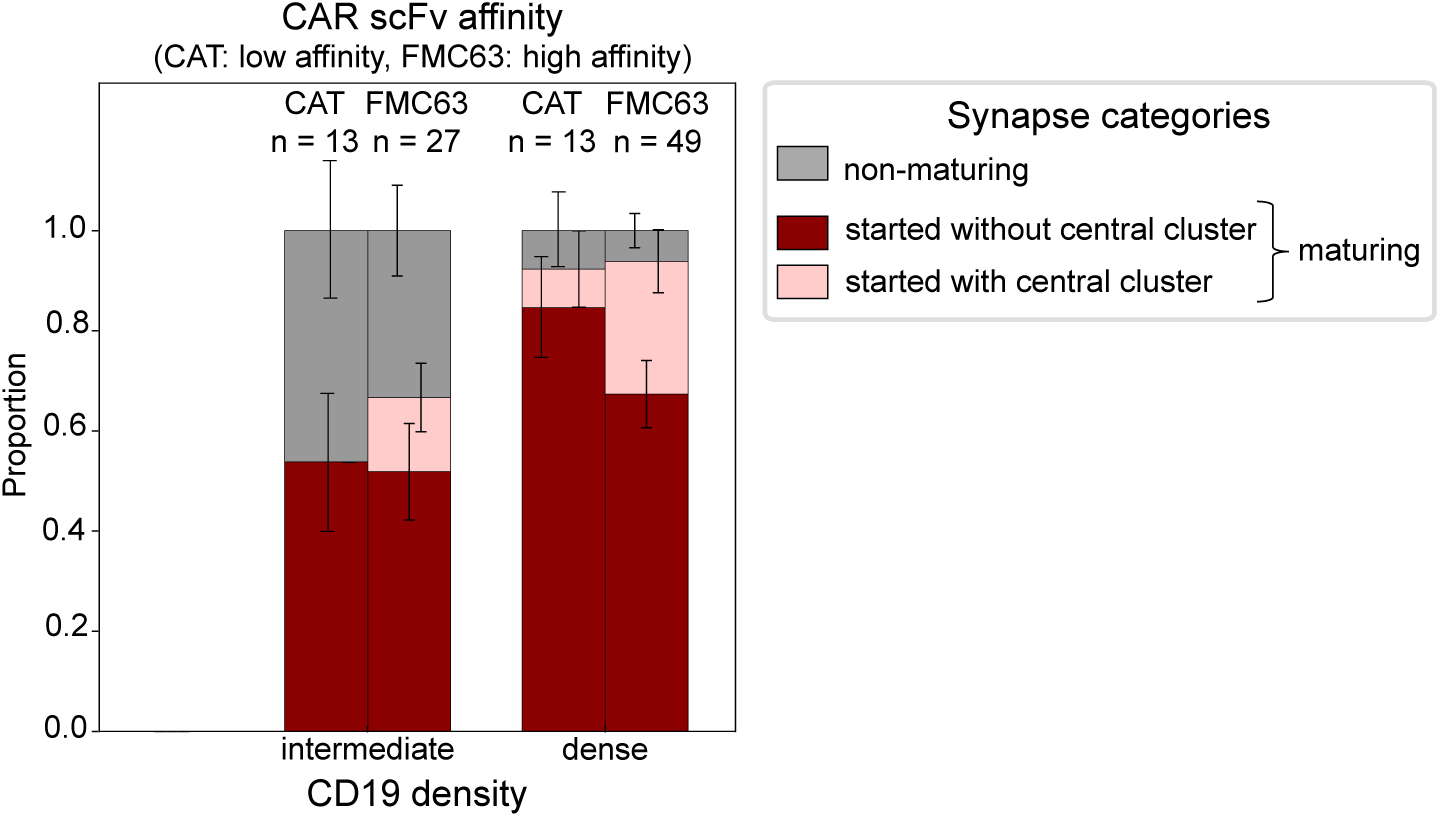
Synapse maturation of high-CAR-expressing CAR-T cells across CD19 densities. Stacked bar plots showing the proportion of high-CAR-expressing cells classified into different synapse maturation categories at intermediate and dense CD19 densities on SLBs for low-affinity (CAT) and high-affinity (FMC63) CAR-T cells. Error bars represent the standard errors of the binomial proportions. Numbers of analyzed cells were: CAT CAR-T cells, *n* = 13 (intermediate) and *n* = 13 (dense); FMC63 CAR-T cells, *n* = 27 (intermediate) and *n* = 49 (dense).

**Figure S5.**
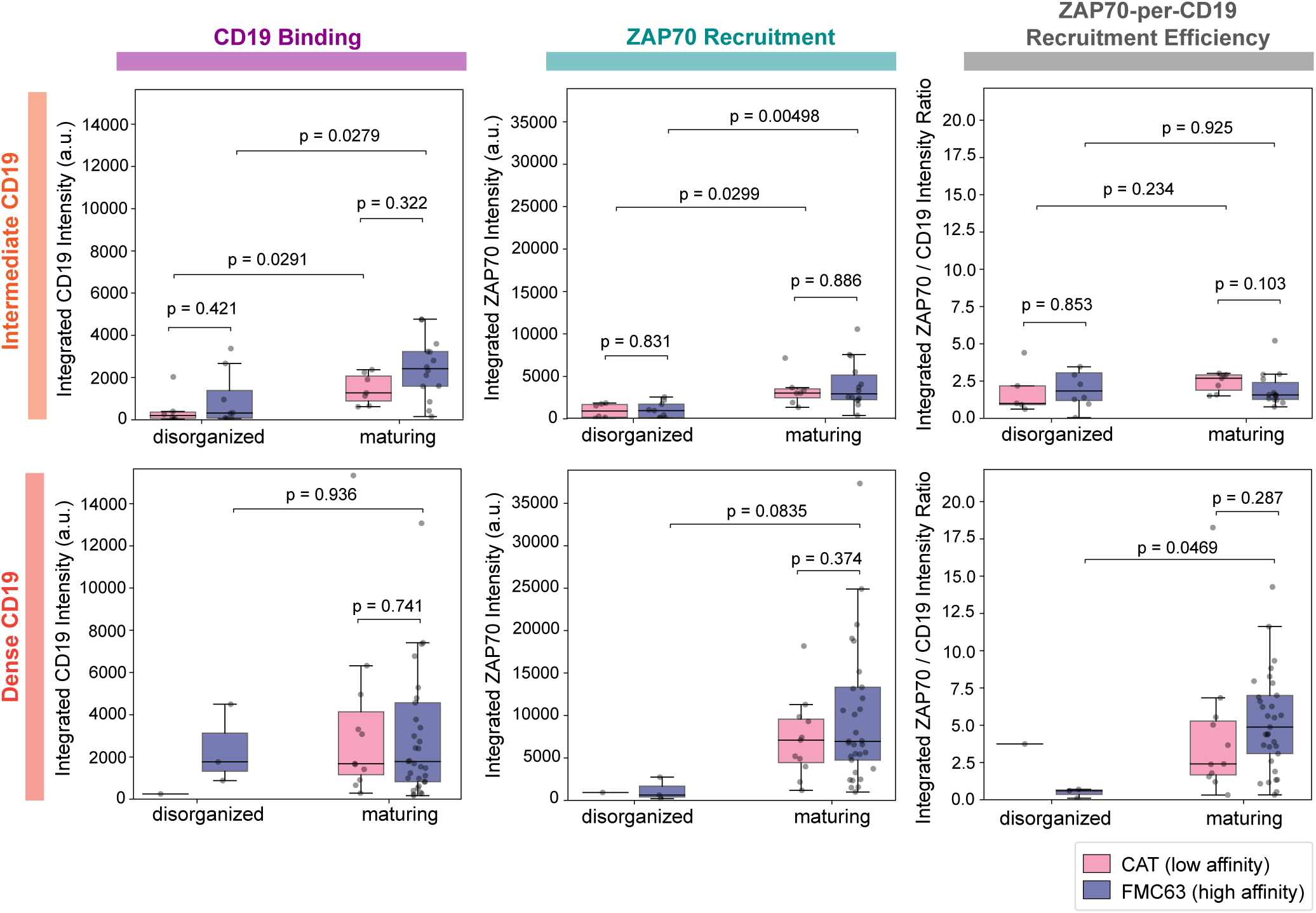
CD19 binding and ZAP70 recruitment in disorganized and maturing synapses across CD19 densities for high-CAR-expressing CAR-T cells. Box plots of the per-cell integrated CD19-JF646 intensity, ZAP70-eGFP intensity, and ZAP70/CD19 intensity ratio for all CAR clusters detected during time-lapse TIRF imaging of disorganized and maturing synapses in low-affinity (CAT) and high-affinity (FMC63) CAR-T cells expressing high CAR levels on intermediate and dense CD19 SLBs. Boxes indicate the median and interquartile range, and individual points represent a single cell. Intermediate CD19: disorganized CAT (*n* = 6) and FMC63 (*n* = 8); maturing CAT (*n* = 7) and FMC63 (*n* = 14). Dense CD19: disorganized CAT (*n* = 1) and FMC63 (*n* = 3); maturing CAT (*n* = 11) and FMC63 (*n* = 33).

**Figure S6.**
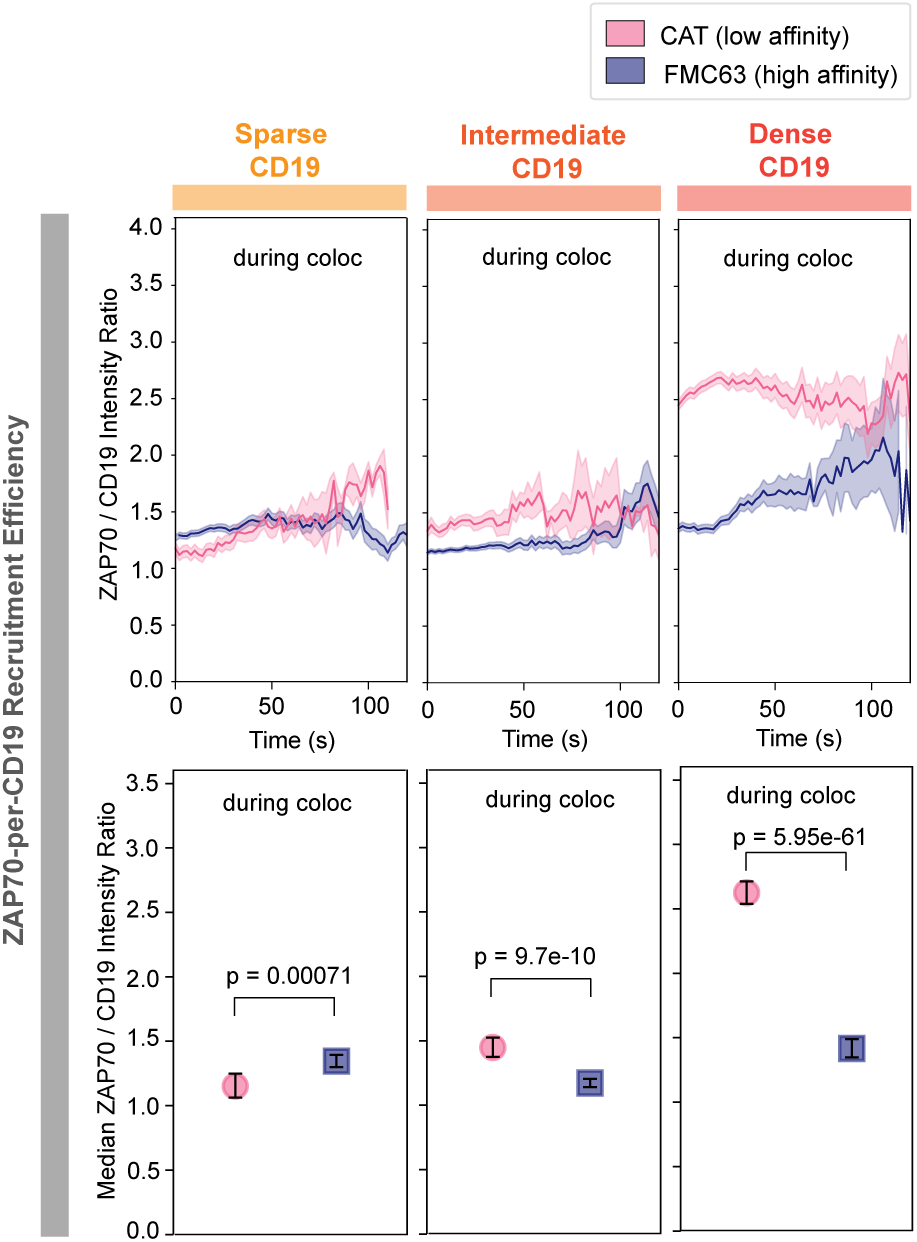
Quantitative analysis of ZAP70 recruitment efficiency in low-CAR-expressing cells that formed maturing synapses. ZAP70/CD19 fluorescence intensity ratio traces, shown as geometric means with SEM estimated from log-transformed values, from small CD19-CAR and ZAP70 microcluster trajectories, together with dot plots showing the corresponding median fluorescence intensities ratio with bootstrap confidence intervals, generated by low-affinity (CAT) and high-affinity (FMC63) CAR-T cells expressing low CAR levels interacting with sparse, intermediate, or dense CD19 SLBs. Number of analyzed tracks under different CD19 densities: sparse CD19, CAT (n = 55) and FMC63 (n = 218); intermediate CD19, CAT (n = 29) and FMC63 (n = 784); dense CD19, CAT (n = 447) and FMC63 (n = 147).

**Figure S7.**
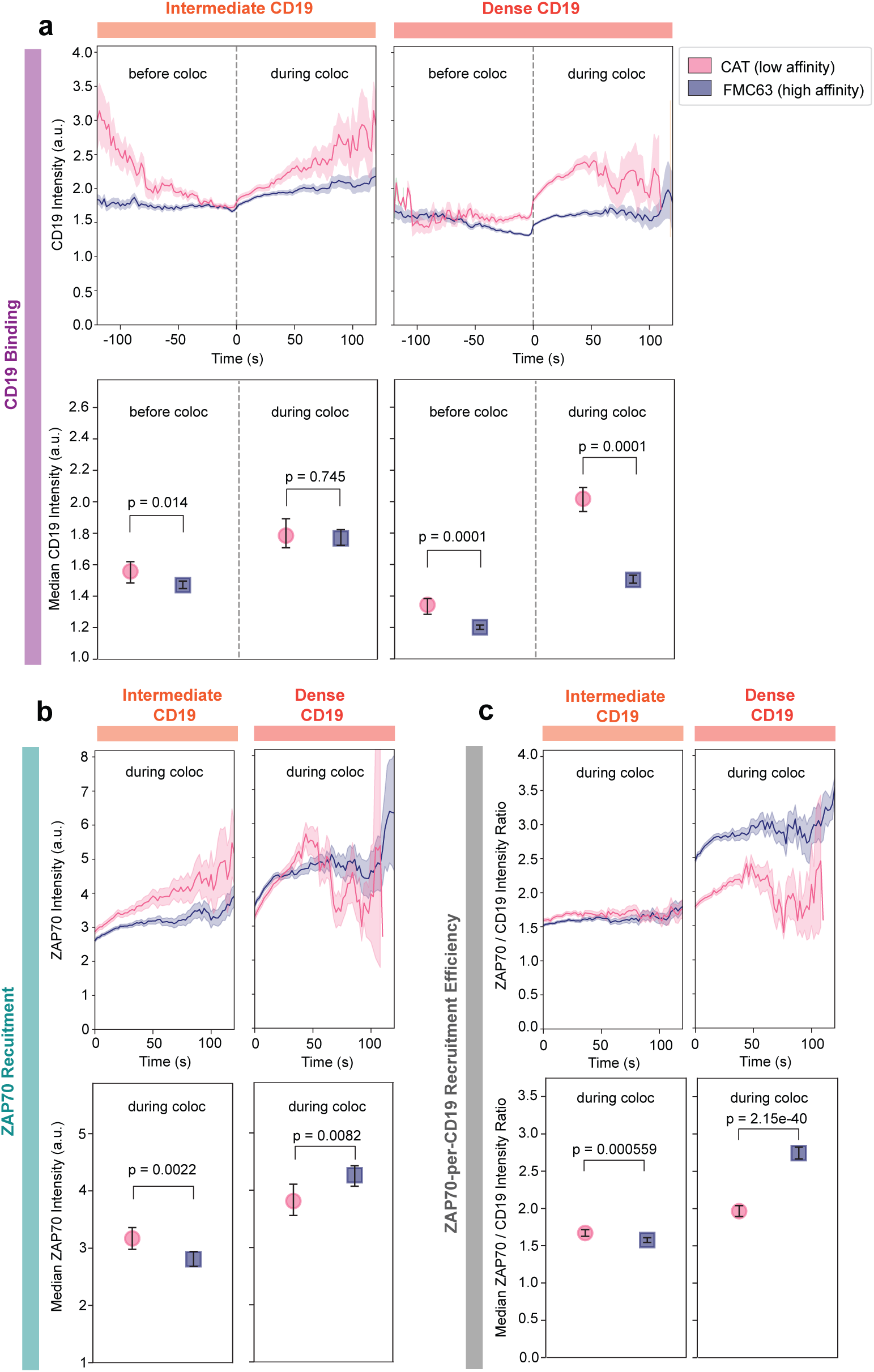
Quantitative analysis of CD19 binding, ZAP70 recruitment, and ZAP70 recruitment efficiency per CD19 molecule in small CAR microclusters before and during ZAP70 recruitment within maturing immunological synapses formed by high-CAR-expressing low-affinity (CAT) and high-affinity (FMC63) CAR-T cells interacting with intermediate and dense CD19 SLBs. Fluorescence intensity traces of **(a)** CD19-JF646, **(b)** ZAP70-eGFP, and **(c)** ZAP70/CD19 ratio, shown as geometric means with SEM estimated from log-transformed values, from small CD19-CAR and ZAP70 microcluster trajectories, together with dot plots showing the corresponding median fluorescence intensities with bootstrap confidence intervals, generated by low-affinity (CAT) and high-affinity (FMC63) CAR-T cells expressing high CAR levels interacting with intermediate or dense CD19 SLBs. Number of analyzed tracks under different CD19 densities: intermediate CD19, CAT (n = 297) and FMC63 (n = 801); dense CD19, CAT (n = 422) and FMC63 (n = 1025).

**Figure S8.**
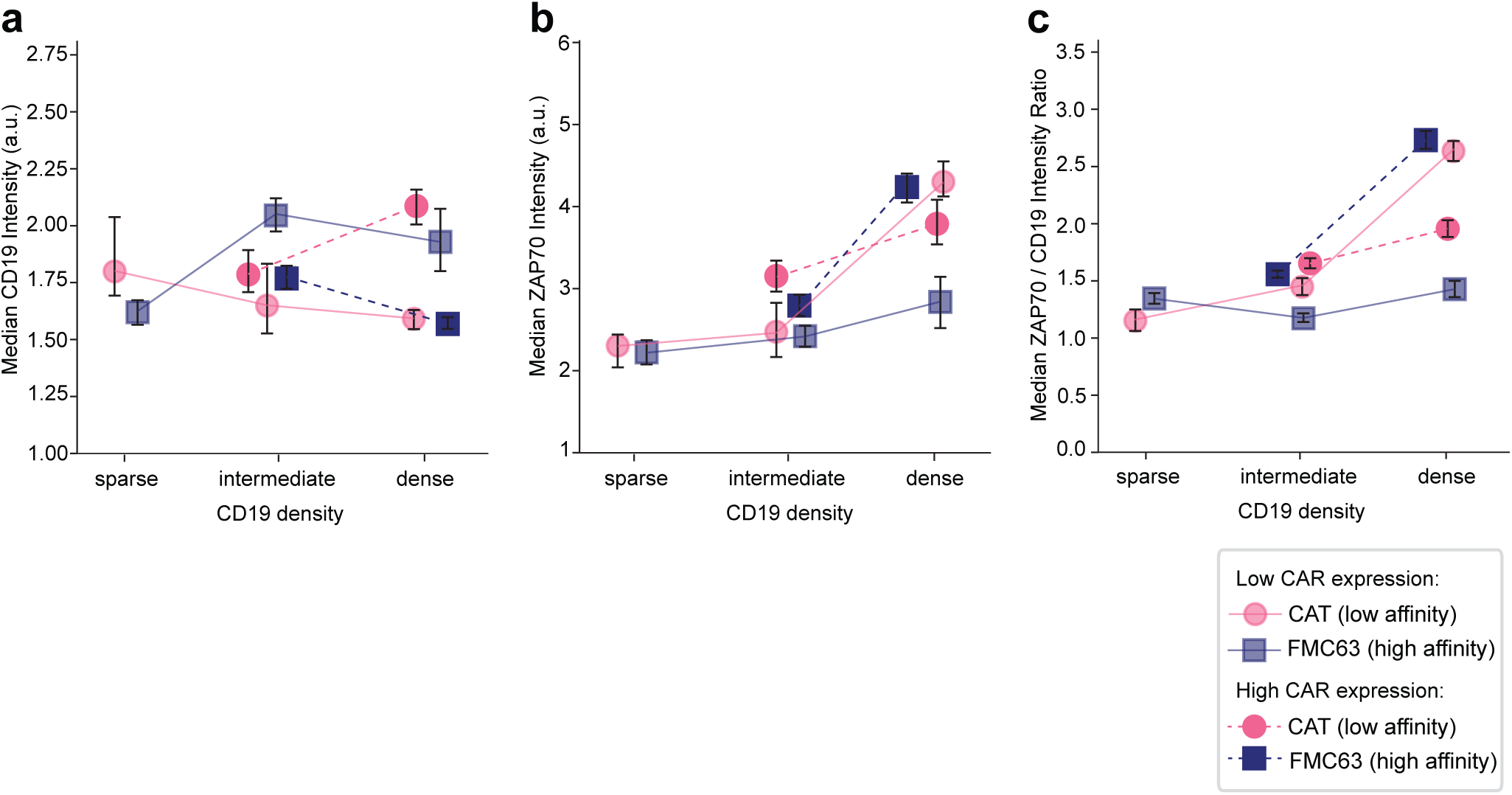
Combined quantitative analysis of CD19 binding, ZAP70 recruitment, and ZAP70 recruitment efficiency in small, maturing CAR microclusters from both low- and high-CAR-expressing T cells interacting with SLBs presenting sparse, intermediate, or dense CD19 densities. **(a)** Median CD19-JF646 fluorescence intensity of small CD19-CAR microclusters during ZAP70 colocalization. Values were reconstructed from the median intensity dot plots in Figure 3b for low-CAR-expressing cells (similar to Figure 3c) and Figure S7a for high-CAR-expressing cells. **(b)** Median ZAP70-eGFP fluorescence intensity of ZAP70 microclusters colocalized with small CD19-CAR microclusters. Values were reconstructed from the median intensity dot plots in Figure 3b for low-CAR-expressing cells (similar to Figure 3d) and Figure S7b for high-CAR-expressing cells. **(c)** Median ZAP70 recruitment efficiency, quantified as the ZAP70-eGFP/CD19-JF646 fluorescence intensity ratio in small CAR microclusters. Values were reconstructed from the median intensity dot plots in Figure S6 for low-CAR-expressing cells (similar to Figure 3e), and from Figure S7c for high-CAR-expressing cells. Number of analyzed tracks for low-CAR-expressing cells under different CD19 densities: sparse CD19, CAT (n = 55) and FMC63 (n = 218); intermediate CD19, CAT (n = 29) and FMC63 (n = 784); dense CD19, CAT (n = 447) and FMC63 (n = 147). Number of analyzed tracks for high-CAR-expressing cells under different CD19 densities: intermediate CD19, CAT (n = 297) and FMC63 (n = 801); dense CD19, CAT (n = 422) and FMC63 (n = 1025).

**Figure S9.**
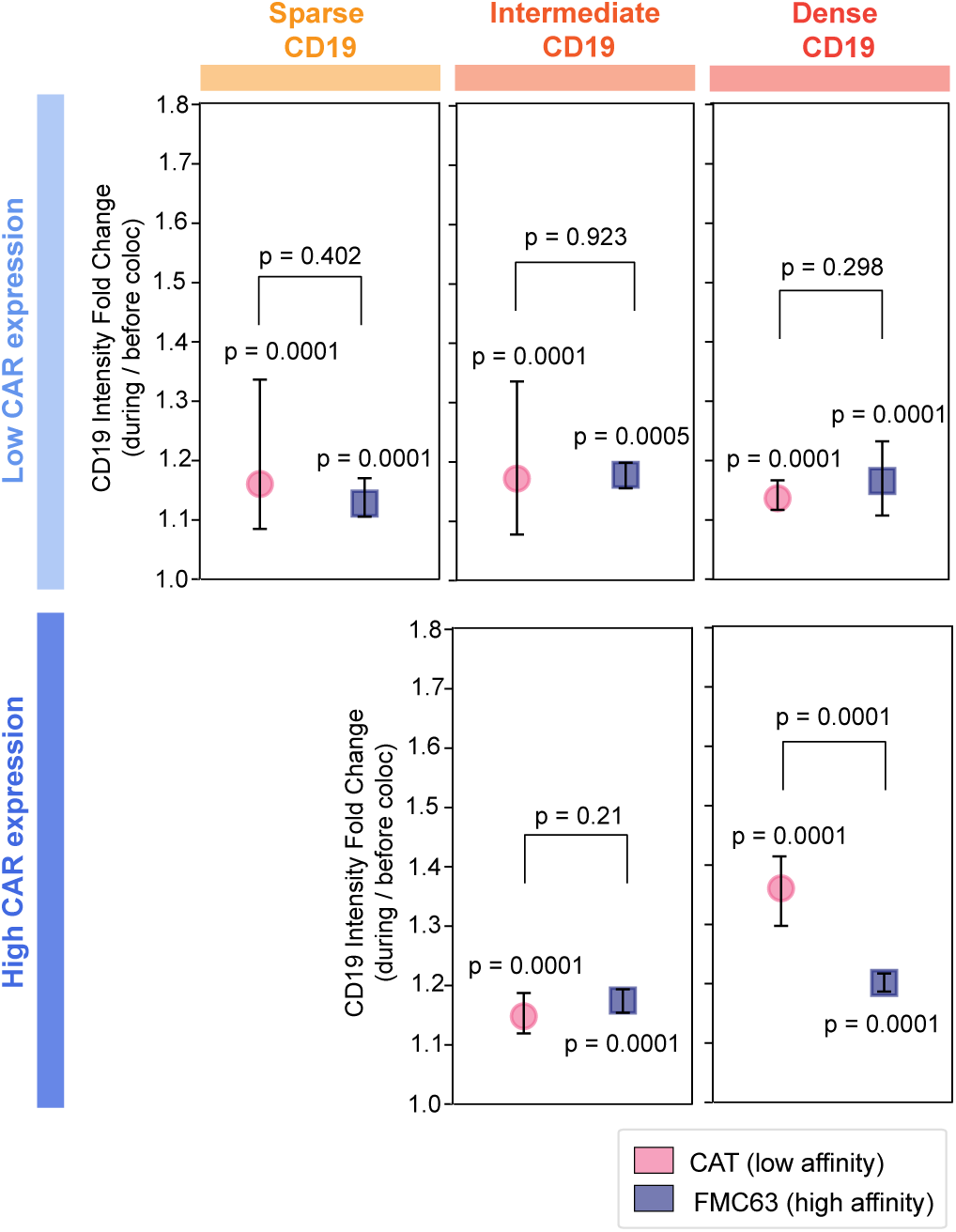
CD19 binding fold change associated with ZAP70 recruitment to small, mature CD19-CAR microclusters in low- and high-CAR-expressing cells across different CD19 densities on SLBs. Dot plots showing the median CD19-JF646 fluorescence intensty fold change with bootstrap confidence intervals. Fold change was calculated as the ratio of median CD19-JF646 fluorescence intensity during ZAP70 colocalization over that before ZAP70 colocalization (values shown in Figure 3b) for each microcluster. Number of analyzed tracks for low-CAR-expressing cells under different CD19 densities: sparse CD19, CAT (n = 55) and FMC63 (n = 218); intermediate CD19, CAT (n = 29) and FMC63 (n = 784); dense CD19, CAT (n = 447) and FMC63 (n = 147). Number of analyzed tracks for high-CAR-expressing cells under different CD19 densities: intermediate CD19, CAT (n = 297) and FMC63 (n = 801); dense CD19, CAT (n = 422) and FMC63 (n = 1025).

**Figure S10.**
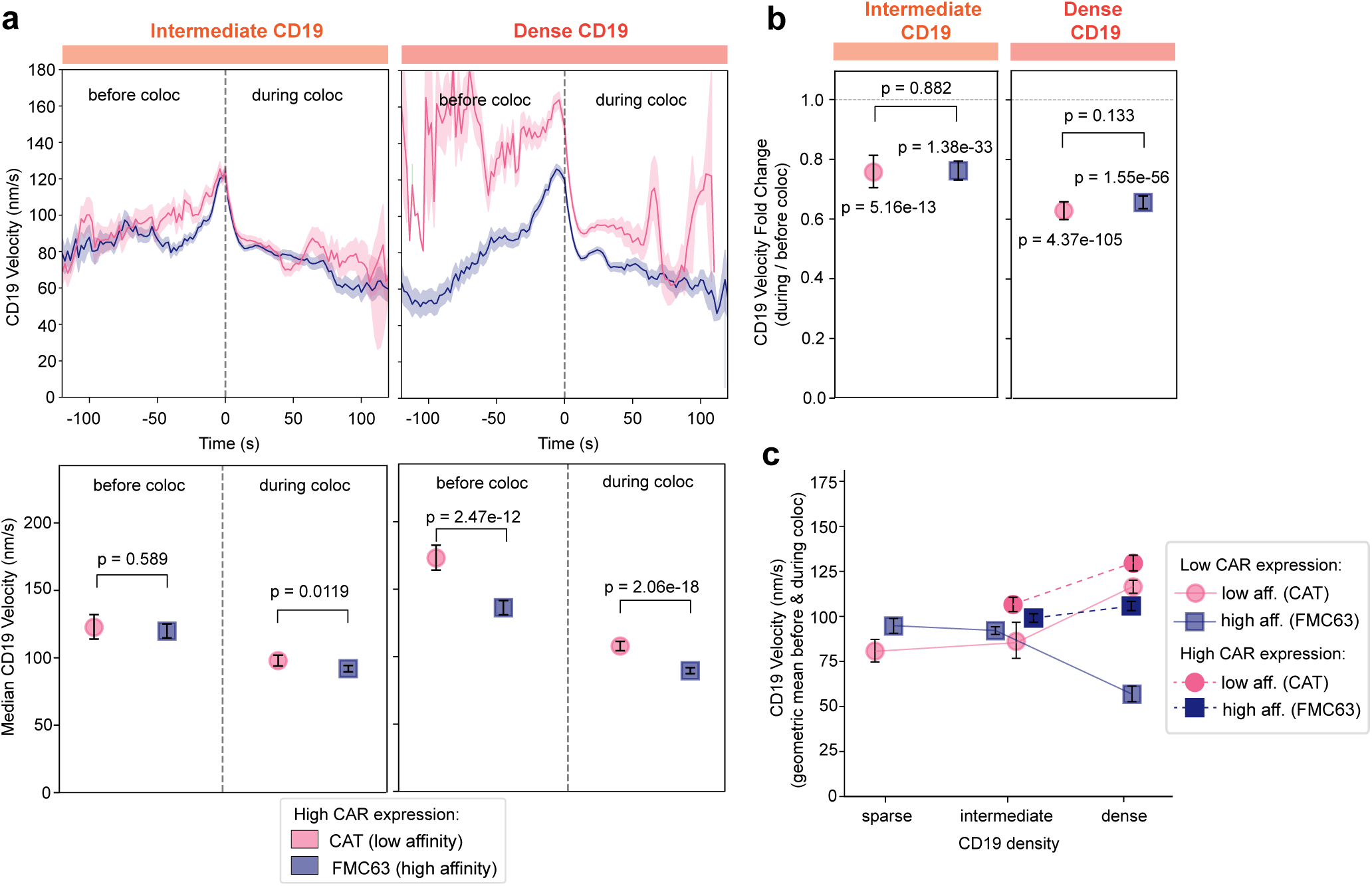
Quantitative analysis of the velocity of small CD19-CAR microclusters before and during ZAP70 recruitment in maturing immunological synapses formed by high-CAR-expressing low-affinity (CAT) and high-affinity (FMC63) CAR-T cells interacting with intermediate and dense CD19 SLBs. **(a)** CD19-JF646 velocity traces, shown as geometric means with SEM estimated from log-transformed values, from small CD19-CAR microcluster trajectories before and during colocalization with ZAP70, together with dot plots showing the corresponding median velocities with 95% nonparametric bootstrap confidence intervals, generated by low-affinity (CAT) and high-affinity (FMC63) CAR-T cells expressing high CAR levels interacting with intermediate or dense CD19 SLBs. **(b)** Dot plots showing the median CD19-JF646 velocity fold change, which was associated with ZAP70 recruitment to small, mature CD19-CAR microclusters in high-CAR-expressing cells, with bootstrap confidence intervals. Fold change was calculated as the ratio of median CD19-JF646 velocity during ZAP70 colocalization over that before ZAP70 colocalization (values shown in Figure S10a) for each microcluster. **(c)** Mean velocity of CD19-CAR microclusters across the full trajectory, including periods before and during ZAP70 colocalization. Values represent the geometric mean of the instantaneous CD19-JF646 microcluster velocities, with 95% nonparametric bootstrap confidence intervals. Values were reconstructed from the median velocity dot plots in Figure 4b for low-CAR-expressing cells (similar to Figure 4c) and Figure S10a for high-CAR-expressing cells. Number of analyzed tracks for low-CAR-expressing cells under different CD19 densities: sparse CD19, CAT (n = 55) and FMC63 (n = 218); intermediate CD19, CAT (n = 29) and FMC63 (n = 784); dense CD19, CAT (n = 447) and FMC63 (n = 147). Number of analyzed tracks for high-CAR-expressing cells under different CD19 densities: intermediate CD19, CAT (n = 297) and FMC63 (n = 801); dense CD19, CAT (n = 422) and FMC63 (n = 1025).

**Figure S11.**
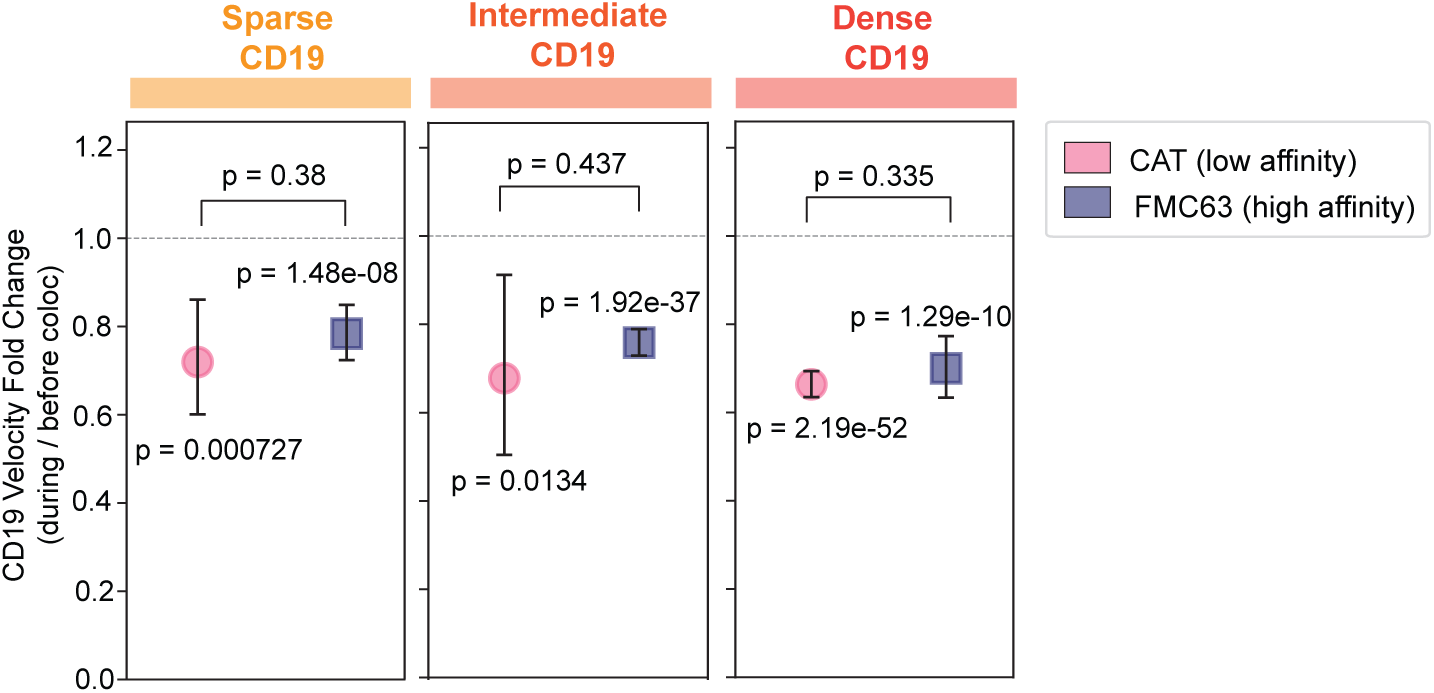
Velocity fold change of small, mature CD19-CAR microclusters associated with ZAP70 recruitment in low-CAR-expressing cells across different CD19 densities on SLBs. Dot plots showing the median CD19-JF646 velocity fold change, which was associated with ZAP70 recruitment to small, mature CD19-CAR microclusters in low-CAR-expressing cells, with bootstrap confidence intervals. Fold change was calculated as the ratio of median CD19-JF646 velocity during ZAP70 colocalization over that before ZAP70 colocalization (values shown in Figure 4b) for each microcluster. Number of analyzed tracks under different CD19 densities: sparse CD19, CAT (n = 55) and FMC63 (n = 218); intermediate CD19, CAT (n = 29) and FMC63 (n = 784); dense CD19, CAT (n = 447) and FMC63 (n = 147).

**Figure S12.**
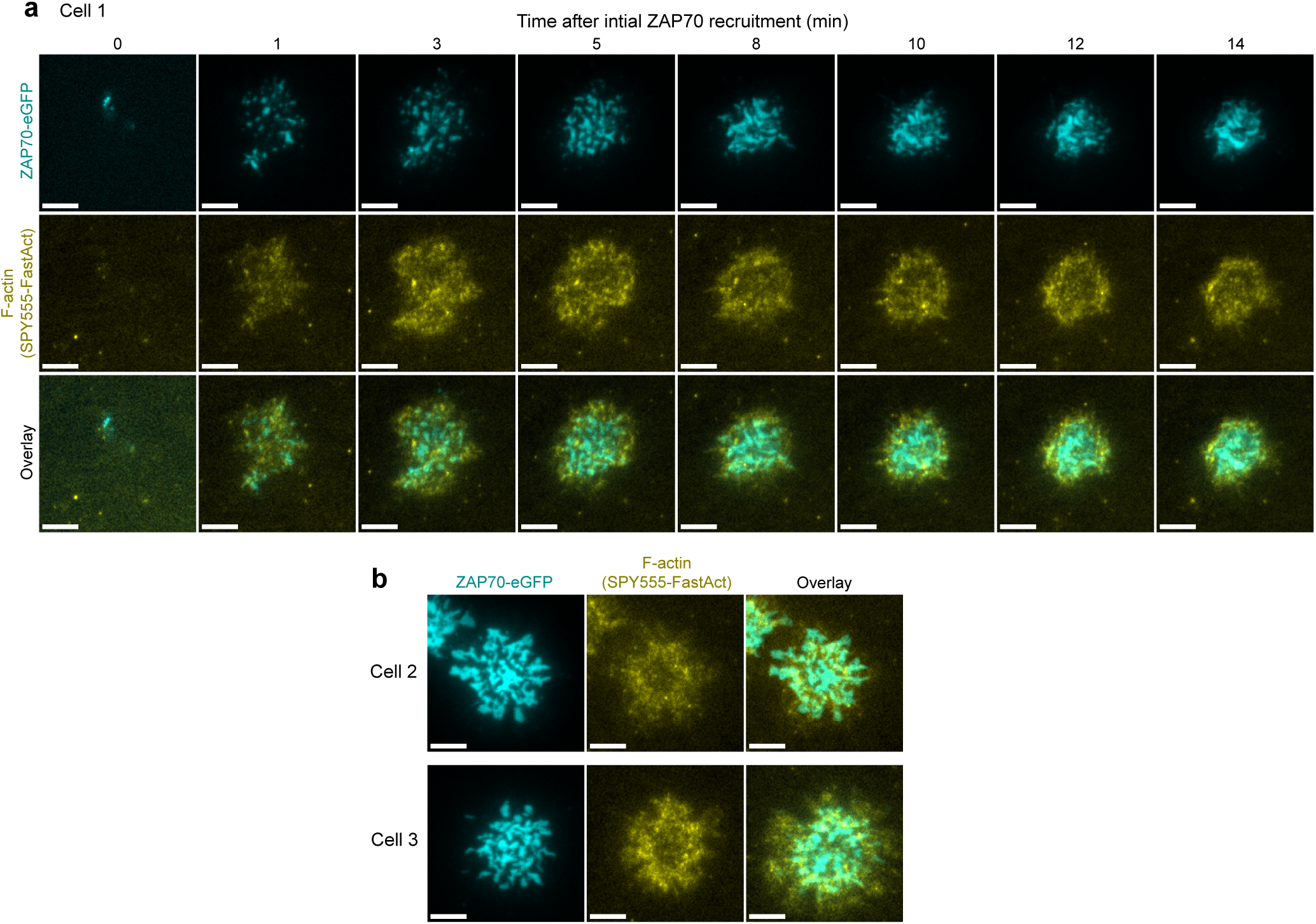
Spatiotemporal organization of F-actin and ZAP70 at CAR immunological synapses reveals that actin dynamics drive the centripetal transport of CAR microclusters. **(a)** Representative TIRF time-lapse images of a single CAR-T cell from initial contact with a CD19-presenting SLB through 14 min of synapse maturation, showing ZAP70 recruitment, formation of a peripheral F-actin-rich ring, and centripetal movement of CAR clusters driven by retrograde F-actin flow. Scale bars are 5 µm. **(b)** Representative TIRF images of two additional CAR-T cells exhibiting a similar spatiotemporal organization as shown in (a), with ZAP70 clusters localized within the peripheral F-actin-rich zone, demonstrating the reproducible association between peripheral actin structures and the centripetal transport of signaling microclusters. Scale bars are 5 µm.

**Figure S13.**
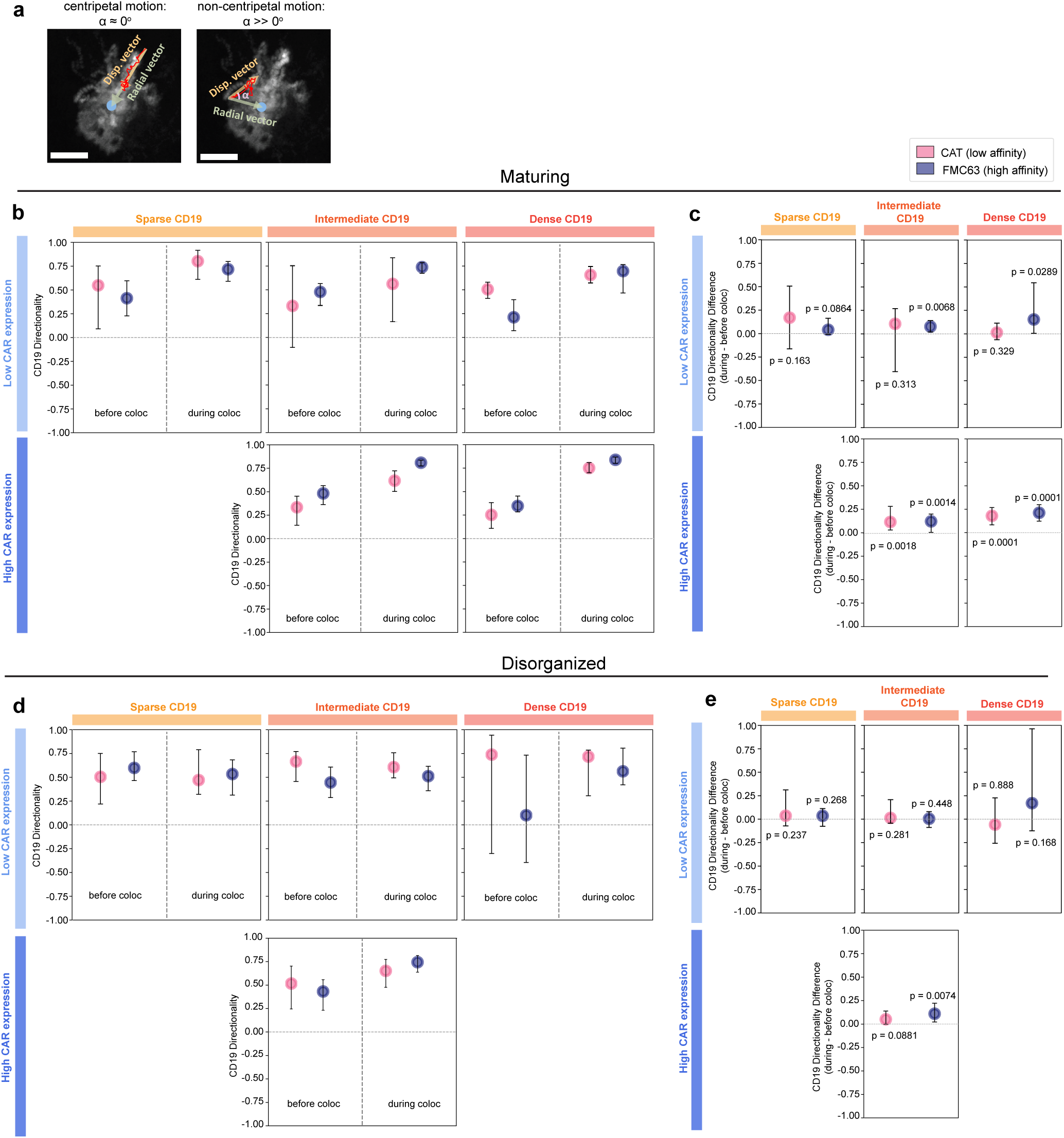
Quantitative analysis of the directionality of small CD19-CAR microclusters before and during ZAP70 recruitment within maturing and disorganized immunological synapses formed by low- and high-CAR-expressing low-affinity (CAT) and high-affinity (FMC63) CAR-T cells interacting with SLBs presenting different CD19 densities. **(a)** Representative TIRF images illustrating quantification of small CD19-CAR microcluster directionality through tracking of the CD19-JF646 fluorescence signal. Directionality was defined by the angle (*α*) between the radial and displacement vector of the microcluster. Values of *α* approaching 0*^◦^* indicate highly centripetal movement, whereas increasing angles reflect progressively less centripetal trajectories. Scale bars are 5 µm. **(b)** Dot plots showing the directionality of individual small CD19-CAR microcluster trajectories in cells forming maturing synapses before and during ZAP70 colocalization, presented as the median with 95% nonparametric bootstrap confidence intervals. Data are expressed as the cosine of *α*, ranging from -1 to +1, where values approaching +1 indicate strongly centripetal movement. **(c)** Dot plots showing the change in directionality of small CD19-CAR microclusters in maturing synapses, quantified as the per-trajectory difference in cos(*α*) during versus before ZAP70 colocalization. A one-sided paired sign-flip permutation test was used to assess whether the median difference exceeded zero. **(d)** Dot plots showing the directionality of individual small CD19-CAR microcluster trajectories in cells forming disorganized synapses before and during ZAP70 colocalization, presented as the median with 95% nonparametric bootstrap confidence intervals. Data are expressed as the cosine of *α*, ranging from -1 to +1, where values approaching +1 indicate strongly centripetal movement. **(e)** Dot plots showing the change in directionality of small CD19-CAR microclusters in disorganized synapses, quantified as the per-trajectory difference in cos(*α*) during versus before ZAP70 colocalization. A one-sided paired sign-flip permutation test was used to assess whether the median difference exceeded zero. Number of analyzed tracks for low-CAR-expressing cells with maturing synapses under different CD19 densities: sparse CD19, CAT (n = 55) and FMC63 (n = 218); intermediate CD19, CAT (n = 29) and FMC63 (n = 784); dense CD19, CAT (n = 447) and FMC63 (n = 147). Number of analyzed tracks for high-CAR-expressing cells with maturing synapses under different CD19 densities: intermediate CD19, CAT (n = 297) and FMC63 (n = 801); dense CD19, CAT (n = 422) and FMC63 (n = 1025). Number of analyzed tracks for low-CAR-expressing cells with disorganized synapses under different CD19 densities: sparse CD19, CAT (n = 106) and FMC63 (n = 308); intermediate CD19, CAT (n = 127) and FMC63 (n = 359); dense CD19, CAT (n = 12) and FMC63 (n = 68). Number of analyzed tracks for high-CAR-expressing cells with disorganized synapses under different CD19 densities: intermediate CD19, CAT (n = 250) and FMC63 (n = 261); dense CD19, CAT (n = 43) and FMC63 (n = 7).

**Figure S14.**
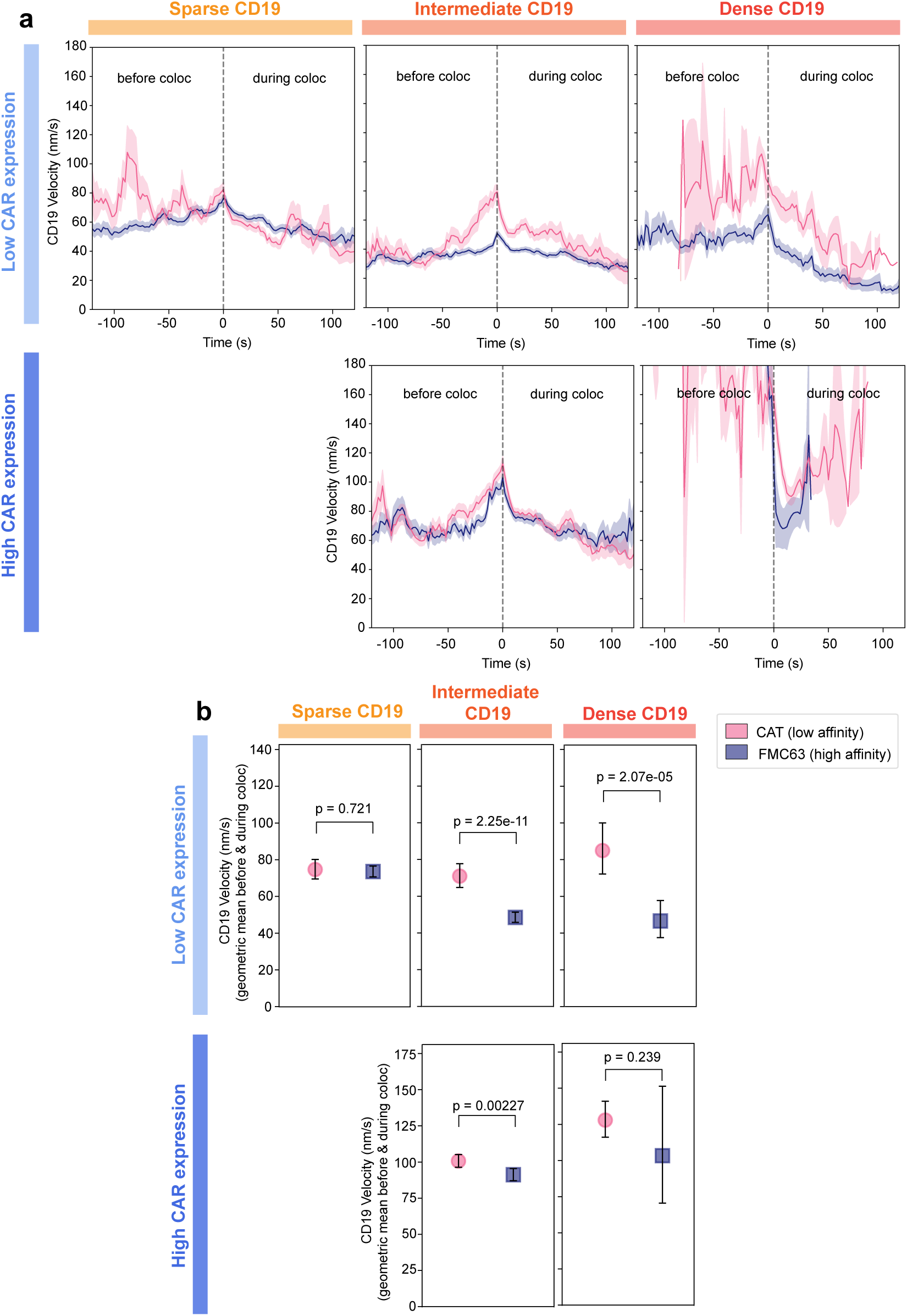
Quantitative analysis of the velocity of small CD19-CAR microclusters before and during ZAP70 recruitment in disorganized immunological synapses formed by low- and high-CAR-expressing low-affinity (CAT) and high-affinity (FMC63) CAR-T cells interacting with different CD19 densities on SLBs. **(a)** CD19-JF646 velocity traces of small CD19-CAR microclusters before and during ZAP70 colocalization, shown as geometric means with SEM estimated from log-transformed values, together with **(b)** dot plots of the corresponding geometric mean velocities with 95% nonparametric bootstrap confidence intervals, from CAR-T cells interacting with CD19-presenting SLBs. Number of analyzed tracks for low-CAR-expressing cells with disorganized synapses under different CD19 densities: sparse CD19, CAT (n = 106) and FMC63 (n = 308); intermediate CD19, CAT (n = 127) and FMC63 (n = 359); dense CD19, CAT (n = 12) and FMC63 (n = 68). Number of analyzed tracks for high-CAR-expressing cells with disorganized synapses under different CD19 densities: intermediate CD19, CAT (n = 250) and FMC63 (n = 261); dense CD19, CAT (n = 43) and FMC63 (n = 7).

**Figure S15.**
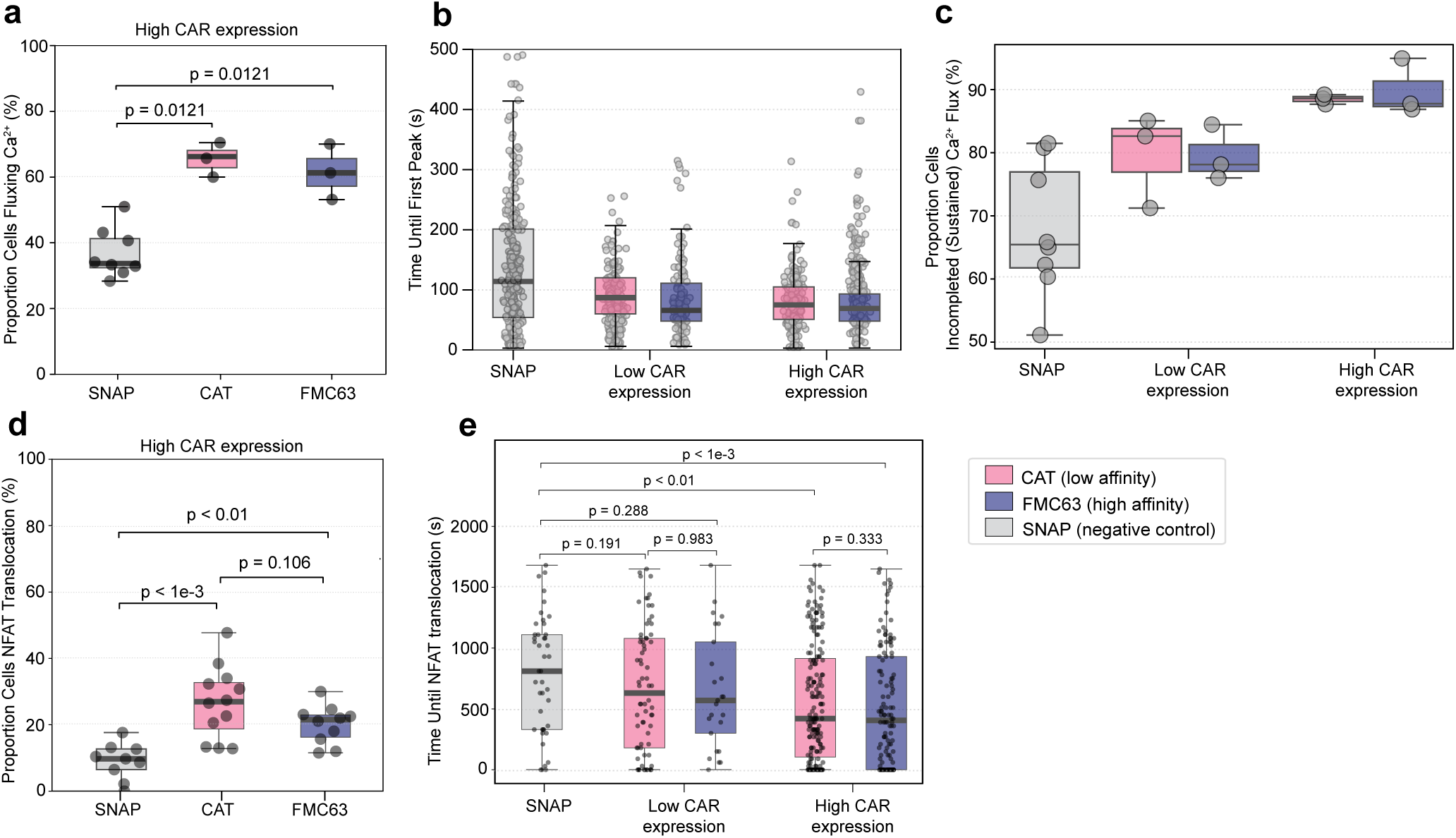
Intracellular Ca^2+^ signaling and NFAT nuclear translocation following low-affinity (CAT) and high-affinity (FMC63) CAR-T cells interacting with dense CD19 or SNAP-functionalized (negative control) SLBs. **(a)** Box plot showing the proportion (%) of high-CAR-expressing low-affinity (CAT) and high-affinity (FMC63) CAR-T cells exhibiting robust intracellular Ca^2+^ flux upon interaction with dense CD19 SLBs or with non-reactive SNAP-functionalized SLBs lacking CD19. Boxes indicate the median and interquartile range, and individual points represent independent replicates. Cell numbers: CAT high expression (n = 293 cells, 3 replicates) and FMC63 high expression (n = 487 cells, 3 replicates) on CD19 SLBs; SNAP control, combined CAT and FMC63 CAR-T cells (n = 654 cells, 8 replicates). **(b)** Box plot showing time duration (s) until the first robust Ca^2+^ peak detected in low- and high-CAR-expressing low-affinity (CAT) and high-affinity (FMC63) CAR-T cells upon interaction with dense CD19 or with non-reactive SNAP SLBs. Boxes indicate the median and interquartile range, and individual points represent independent cells. Cell numbers: CAT low expression (n = 245 cells, 3 replicates), FMC63 low expression (n = 121 cells, 3 replicates), CAT high expression (n = 231 cells, 3 replicates), FMC63 high expression (n = 302 cells, 3 replicates); SNAP control, combined CAT and FMC63 CAR-T cells (n = 283 cells, 8 replicates). **(c)** Box plot showing the proportion (%) of low- and high-CAR-expressing low-affinity (CAT) and high-affinity (FMC63) CAR-T cells exhibiting sustained (incompleted) Ca^2+^ flux upon interaction with dense CD19 or with non-reactive SNAP SLBs. Boxes indicate the median and interquartile range, and individual points represent independent replicates. Cell numbers: CAT low expression (n = 245 cells, 3 replicates), FMC63 low expression (n = 121 cells, 3 replicates), CAT high expression (n = 231 cells, 3 replicates), FMC63 high expression (n = 302 cells, 3 replicates); SNAP control, combined CAT and FMC63 CAR-T cells (n = 283 cells, 8 replicates). **(d)** Box plot showing the proportion (%) of high-CAR-expressing low-affinity (CAT) and high-affinity (FMC63) CAR-T cells exhibiting robust NFAT nuclear translocation upon interaction with dense CD19 or with non-reactive SNAP SLBs. Boxes indicate the median and interquartile range, and individual points represent independent replicates. Cell numbers: CAT high expression (n = 179 cells, 12 replicates), FMC63 high expression (n = 126 cells, 10 replicates); SNAP control, combined CAT and FMC63 CAR-T cells (n = 43 cells, 9 replicates). **(e)** Box plot showing the time duration (s) until a robust NFAT nuclear translocation detected in low- and high-CAR-expressing low-affinity (CAT) and high-affinity (FMC63) CAR-T cells interacting with dense CD19 or with non-reactive SNAP SLBs. Boxes indicate the median and interquartile range, and individual points represent independent cells. Cell numbers: CAT low expression (n = 70 cells, 5 replicates), FMC63 low expression (n = 25 cells, 5 replicates), CAT high expression (n = 179 cells, 12 replicates), FMC63 high expression (n = 126 cells, 10 replicates); SNAP control, combined CAT and FMC63 CAR-T cells (n = 43 cells, 9 replicates).

